# Osmolyte structural and thermodynamic effects across the protein folding landscape

**DOI:** 10.1101/2025.02.07.637036

**Authors:** Ander Francisco Pereira, Leandro Martínez

**Affiliations:** Institute of Chemistry and Center for Computing in Engineering & Science, Universidade Estadual de Campinas (UNICAMP), 13083-861 Campinas, SP, Brazil

## Abstract

In this work, the structure and thermodynamics of urea and TMAO preferential interactions across the complete folding landscape of the β-sheet SH3 domain and the helical B domain of protein A (BdpA) are characterized. There is a high correlation between preferential interactions and the surface area of the denatured states, despite the fact that the chemical nature of the exposed surfaces upon denaturation differs from those of native states. For SH3, the denaturation always proceeds with an increase in surface area, such that the qualitative effect of cosolvents on the stability of all denatured states can be inferred from the preferential interactions in the native state. On the other hand, for BdpA, partially denatured states exist that are destabilized by urea; thus, unfolding pathways can be modulated by the cosolvent. Both urea and TMAO form hydrogen bonds with the proteins, which are weakened upon denaturation, and nonspecific interactions, which are strengthened in unfolded structures. Backbone, side chain, and specific residue contributions to distribution functions are obtained, illustrating, for instance, the crucial participation of urea-backbone and nonpolar-cosolvent interactions in the solvation mechanisms. Obtaining these results was possible using a novel computational pipeline to represent solvation structures throughout complete folding landscapes by means of coarse-grained and atomistic simulations and, crucially, the analysis of solvation using minimum-distance distribution functions and the Kirkwood-Buff solvation theory. Cosolvent effects on transfer free energies match experimental data within 1 kcal mol^-1^, supporting the nuanced description of the osmolyte-protein interplay. The proposed methods can be applied to the study of solvation structure and thermodynamics in other complex molecular systems undergoing large conformational variations, such as non-biological macromolecules and aggregates.

## Introduction

Urea and trimethylamine N-oxide (TMAO) are osmolytes that exert different effects on protein stability, with urea typically acting as a denaturant and TMAO as a protectant.^1–6^ These effects on protein stability can be analyzed, at a first level of approximation, from interactions with the protein in its native state: osmolyte exclusion from the protein surface favors compact structures, typically linked to folded forms.^7,8^ Conversely, preferential protein-osmolyte interactions promote the exposure of the protein residues and thus denaturation.^9,10^ These are, in fact, the well-known behaviors of urea and TMAO, providing the foundation for the understanding of their effects on protein stability.^6,10,11^ This rationale is oversimplified for proteins because denaturation alters the overall chemical nature of the protein surface. Therefore, interactions observed in the native state do not fully capture the complex interplay between water and cosolvents in the unfolded ensemble.^6,12–14^

Obtaining preferential interaction parameters for multiple protein conformations allows for quantitative analysis of the impact of cosolvents on folding equilibrium.^10,12,15^ Determining the absolute cosolvent effect on the folding equilibria requires either directly measuring equilibrium constants in both pure solvent and cosolvent solutions or determining the transfer free energies of the native and denatured states.^2^ The experimental characterization of the folding equilibrium is challenging because the conformations of native and unfolded ensembles may differ in different protective and denaturant conditions, and experiments depend on a reduced variable to classify the ensembles: the protein activity, the secondary structure content, a specific NMR signal, etc. From a computational modeling perspective, the challenges are 1) the proper representation of the interactions; 2) the difficulty in sampling diverse macromolecular conformational states; and 3) the molecular representation of the solvation in structural diverse ensembles and computing observables. In the context of the present work, the force fields developed for reproducing protein preferential interactions of urea^16^ and, more recently, for TMAO,^17^ address the first challenge, and we build our work on top of these developments.

The challenge of representing diverse macromolecular conformations has been tackled with roughly two approaches: simulating small models of peptides or amino acid residues,^17–20^ or through the generation of unfolded states of larger proteins by means of long simulations in denaturing conditions^14^ or with constructs based on experimental data.^13^

The simulations with small protein models allowed the quantitative computation of observables such as preferential interactions and transfer free energies, and provided many essential insights into the mechanisms of urea and TMAO effects on protein stability. In a key study, Montgomery Pettitt and co-workers obtained a detailed characterization of the solvation of a helical decalanine peptide.^18^ They focus on the competing roles of TMAO and urea and were able to show that denatured states of the helix display enhanced preferential solvation by urea. TMAO was also observed to dehydrate the helix, although to a lesser extent, such that the transfer free energies follow the expected trends. They obtain transfer free energies by explicitly computing, using generalized replica exchange simulations, solvation free energies at varying cosolvent concentrations. More recently, Emma-Shea and co-workers revisited the urea-TMAO competing mechanisms on protein stability by modeling peptides in solutions of urea and TMAO.^17^ Notably, they reparametrized the TMAO force-field, obtaining preferential exclusion consistently with experimental data. In parallel, simulations of small peptides in urea showed that urea is preferred over water by all but charged amino acid side chains, supporting that direct hydrophobic interactions are an essential part of the urea-induced denaturation thermodynamics.^21^

To demonstrate that the conclusions obtained on small peptide constructs can be transferred to proteins, simulations of protein denaturation were necessary. The key difference is that, for larger proteins, the chemical nature of the residues exposed upon denaturation differs, on average, from that of the surface of the native state. Effectively, since urea interacts more favorably than water with most side chains,^21^ the exposure of the core of the protein leads to urea early intrusion into the protein globule and progressively enhances van der Waals interactions, as demonstrated by Berne and co-workers.^14^ These observations are confirmed by the simulation of experimentally-supported denatured states of Ubiquitin in concentrated urea solutions.^13^

The missing link between the small-model and large-protein simulations is the ability to comprehensively describe the solvation structure and thermodynamics of denatured states. The computation of protein-cosolvent preferential interaction parameters by means of Kirkwood-Buff theory is a well-developed field.^22,23^ Nevertheless, obtaining both a molecular picture of solvation and the solvation thermodynamic parameters for denatured states remains a challenge. A step forward was the demonstration by Martínez and Shimizu that the connection between the solvation shell picture of solute-solvent interactions and the solvation thermodynamics can be obtained for structures of arbitrary shape using minimum-distance distribution functions (MDDFs).^24^ We also developed specialized computational analysis tools to leverage this concept,^25^ allowing the study of structurally complex molecular environments with thermodynamic rigor.^12,26^

Given that MDDFs can be used to characterize the solvation of denatured states, we here exploit their combination with Structure-Based Models (SBMs)^27^ and atomistic simulations to generate complete folding landscapes of proteins in solvent mixtures. We characterize the solvation structures of two proteins with distinct topologies: the β-sheet SH3 domain^28^ and the α-helical B domain of protein A (BdpA)^29^ in aqueous urea and TMAO, and their preferential interaction coefficients and transfer free energies at the level of individual denatured states.

The contributions of the present study to the current understanding and modeling of osmolyte-protein interactions are three-fold: first, from a mechanistic perspective, we show in unprecedented detail how the molecular interactions vary across the native and denatured protein states, understanding the nuanced competition between water and cosolvent for specific sites and chemical groups of the macromolecules. From a thermodynamic perspective, it is the first, to our knowledge, computation of transfer free energies of atomistic descriptions of denatured ensembles, with implications for the role of cosolvents on protein folding pathways. Third, the computational pipeline herein described can be applied to other complex macromolecular-solvent systems, particularly for leveraging the use of shape-adaptive distribution functions to connect the structure and thermodynamics of solvation.

## Results

### The folding landscape of model proteins

Understanding how cosolvents affect the protein folding free-energy surface requires the characterization of the protein conformational variability. The Energy Landscape (EL) theory^30,31^ provides a framework for modeling protein-folding mechanisms, and forms the basis for simulations using Structure-Based Models (SBMs).^27^ SBMs are demonstrated to provide coarse-grained representations of the protein folding energy surfaces and structural ensembles of model proteins.^32–41^ In this work, SBMs are initially used to sample the conformational landscape of the SH3 (a β-fold model, Figure 1A) and BdpA (a helical domain, Figure 1B).^42^ The folding mechanisms of these models have been extensively characterized by experimental and computational methods, including by SBM simulations.^32–36,42,43^

**Figure 1.**
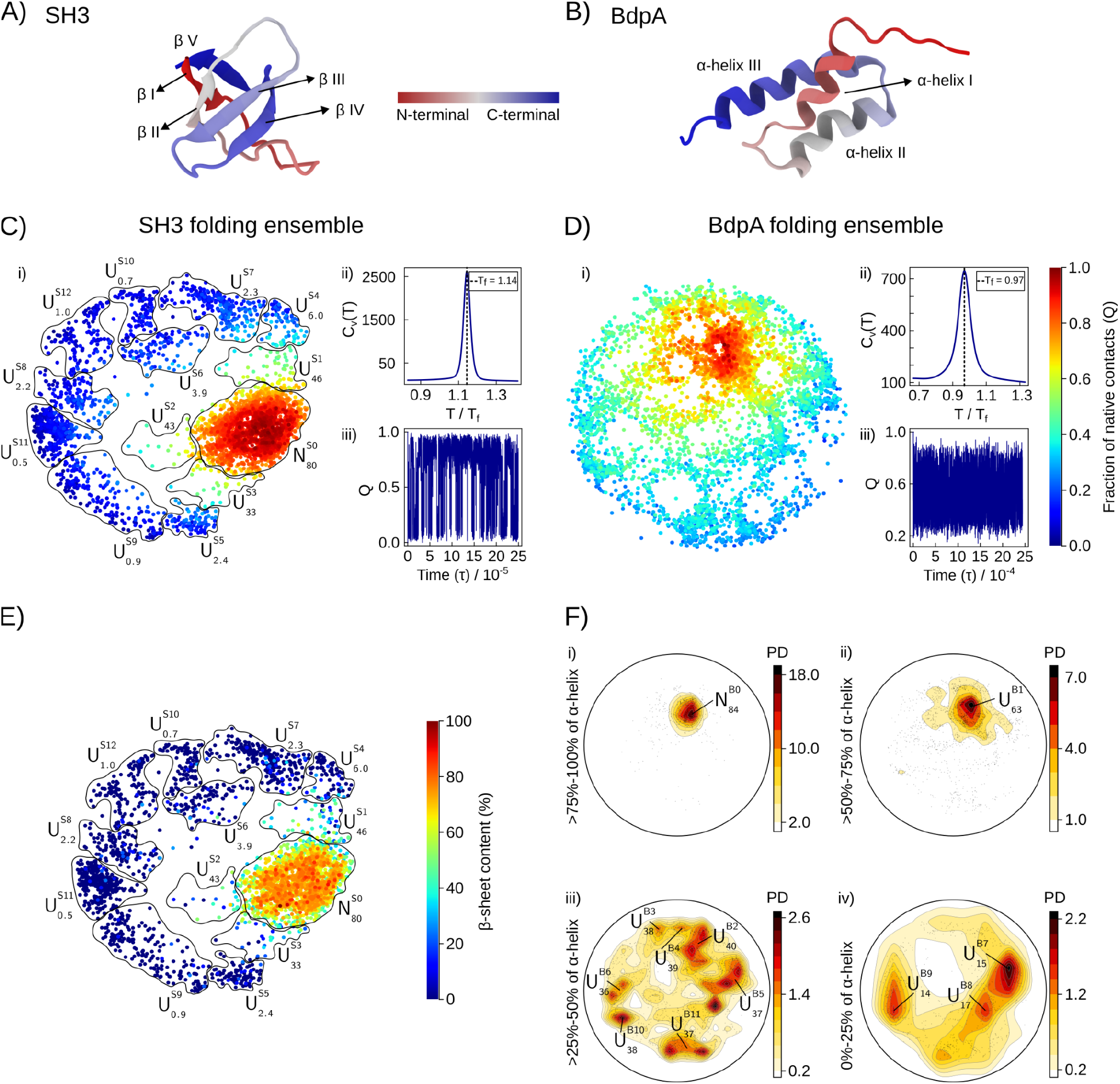
Folding ensembles of SH3 and BpdA. A) and B) Cartoon representation of SH3^28^ and BdpA^29^ native folds. C) and D) SBM folding ensembles characterized by i) the ELViM projection, ii) the specific heat (C_v_) as a function of temperature, and iii) the fraction of native contacts (Q) as a function of time, in reduced units (τ). Each dot in i) represents one structure in a color scale of Q, ranging from 0 (unfolded state) to 1 (folded state). E) Folding projection of SH3 colored by secondary structure content, and definition of the conformational subsets. There is a high correlation between Q (panel C) and the secondary structure content. F) Secondary structure dimension of the 2D projection for BpdA, which is required for classifying unfolded states.^42^ In Figures E) and F), labels are assigned to clusters of native (N) and unfolded (U) states, with a subscript indicating the average secondary structure content and the superscript the order of decreasing fraction of native contacts.

SBM simulations can be performed at the folding temperature of the model, obtaining multiple folding and unfolding events that allow the reconstruction of the folding free-energy surface. For SH3, the overall properties of the SBM sampling are shown in Figure 1C, and the protocol and characterization of the SBM folding are available in Supplementary Information Figures S1 and S2. For BdpA, the characterization of the folding landscapes is shown in Figure 1D, and further details were recently published.^42^ Single sharp peaks in the C_v_(T) profiles (Figures 1C and 1D) indicate two-state folding mechanisms, but the folding transition is sharper for SH3 than for BdpA. In Figures 1C and 1D, panels iii), the exhaustive sampling of native contacts (Q) over time can be observed, an indicator of the proper sampling of the conformational landscapes.

Visualization of the folding landscape requires techniques for dimensionality reduction. Thus, for each SBM simulation, the sampled folds were projected in a two-dimensional space using the *Energy-Landscape Visualization Method* (ELViM).^35,44^ In panels i) of Figures 1C and 1D, each dot represents a conformation obtained in the SBM simulation, and the distances of the dots in the projection are optimized to correlate with the dissimilarity of the models.^35^ The colors of the dots, from blue to red, represent completely unfolded (Q = 0) to completely folded (Q = 1) conformations. In Figures 1C and 1F labels are assigned to clusters of conformations: N and U represent the native basin and unfolded subsets, with a subscript indicating the average secondary structure content of the cluster, and the superscripts *S* or *B* for SH3 or BdpA followed by the classification of the cluster in terms of decreasing Q. For example, 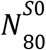 is the native cluster of SH3, which displays on average 80% of the secondary structure of the crystallographic model. Alternatively, 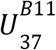 is, in Figure 1F, an unfolded cluster of BdpA with 37% of its native secondary structure content, and the 11th cluster in decreasing amount of native contacts. This labeling will be used along the manuscript to reference the subsets of the folding landscape of the models. Comprehensive structural characterizations of each ensemble are shown in Supplementary Figures S3, S4, and S5, and Table S1.

Figures 1C and 1D show that the folded ensembles (in red) are clustered, as expected from the small native-state conformational variability. On the other hand, unfolded states are spread over the projection, because of their structural diversity. The folding variability of SH3 is captured by the 2D projection: there is a clear native-state basin 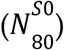, separated from a range of unfolded states 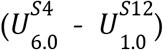 by only three intermediate sets of conformations 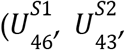, and 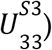. Furthermore, there is a high correlation between the fraction of native contacts (Figure 1C) and the secondary structure content (Figure 1E).

In contrast, the folding space of BdpA displays multiple pathways connecting the native (*N*) and unfolded (*U*) states via structures with similar Q (Figure 1D). Furthermore, partially unfolded states can be found in the native basin, and the secondary structure content needs to be added as an additional dimension to properly differentiate conformations.^42^ Thus, in Figure 1F, projections are split into ranges of protein ellipticity. Panels i), ii), iii), and iv) depict the ensembles within 75-100%, 50-75%, 25-50%, and 0-25% of α-helix content, respectively. The 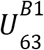 state, in panel ii) of Figure 1F, is particularly interesting: it is a partially denatured state, characterized by the partial loss of the secondary structure of helix I of BdpA, but with mostly preserved native contacts (Supplementary Table S1 and Figures S3 and S5). The denaturation of helix I is widely recognized to be fundamental for the folding equilibrium of BdpA in water.^42,45,46^ The notable interactions of this 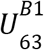 state and its destabilization in the presence of urea will be discussed in detail.

The above characterization of the SH3 and BdpA ensembles is crucial for the primary objective of this work, i.e., to thoroughly investigate urea and TMAO interactions and thermodynamic effects on the protein folding landscapes. With this aim, we obtained all-atom reconstructions of 5000 conformations from the SBM simulations, and each of these atomic models was simulated in aqueous solutions of urea or TMAO at 0.5 mol L^-1^. After equilibration with constrained protein backbone coordinates, the solvated systems were simulated for 10 ns. For each state, as classified in Figure 1, this implied from 210 to 34130 ns of simulations (21 to 3413 structures in each subset), from which the average structure and thermodynamics of the solvents were obtained.

### Osmolyte effects on the folded and unfolded states of SH3

We first focus on the effect of osmolytes urea and TMAO on the folding ensemble of the β-sheet fold of the SH3 domain, which displayed a simpler conformational landscape. Urea is a known denaturant, while TMAO is a stabilizer known to counteract the denaturing effect of urea.^17,47,48^ The structural characterization of protein-solvent interactions was possible using Minimum-Distance Distribution Functions, and the Kirkwood-Buff theory of solvation. The ComplexMixtures.jl^25^ package was developed to perform these analyses. The thermodynamics of protein-solvent interactions was characterized by measuring preferential interaction parameters, the Wyman linkage relation, and transfer free energies.^1,10,49^

MDDFs are distribution functions useful for characterizing the structure of solutions of solutes and solvents of complex shapes,^24^ particularly the unfolded states of proteins.^12^ MDDFs are obtained by computing the minimum-distances between any solute atoms and solvent molecule atoms, and thus retain a clear “solvent-shell” interpretation independently of the complexity of the structures involved. We have shown that a proper normalization of these distributions allows the practical computation of KBIs from the minimum-distance counts,^24^ thus connecting the microscopic picture of solvation to the macroscopic preferential interactions and free-energies of solvation.

Figures 2A-C display the MDDFs of water and urea relative to the protein in the native fold basin 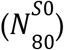 and a selected unfolded state 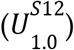, with 2B and 2C being insets of the most prominent peaks of these distributions. The 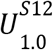 state was chosen for displaying the greatest preferential interactions with urea. Figure 2A shows clear density augmentation of water and urea in the protein vicinity. Both water and urea form hydrogen bonds (H-bonds) with the protein (as evidenced by the peaks at ∼1.9 Å). These H-bond peaks are greater in the native subset (Figure 2B). On the other hand, the MDDFs for the denatured state become greater at distances of nonspecific interactions (>2.5 Å). Thus, water and urea display, in average, stronger H-bonds with the native state, but stronger nonspecific interactions with the denatured states. Both these interactions contribute to urea accumulation around the protein, and as denatured states have a greater surface area (Figure S6), a greater number of protein-solvent interactions is present. The strengthening of the nonspecific interactions upon denaturation is linked to the increased exposure of nonpolar residues (Figure S7).

**Figure 2.**
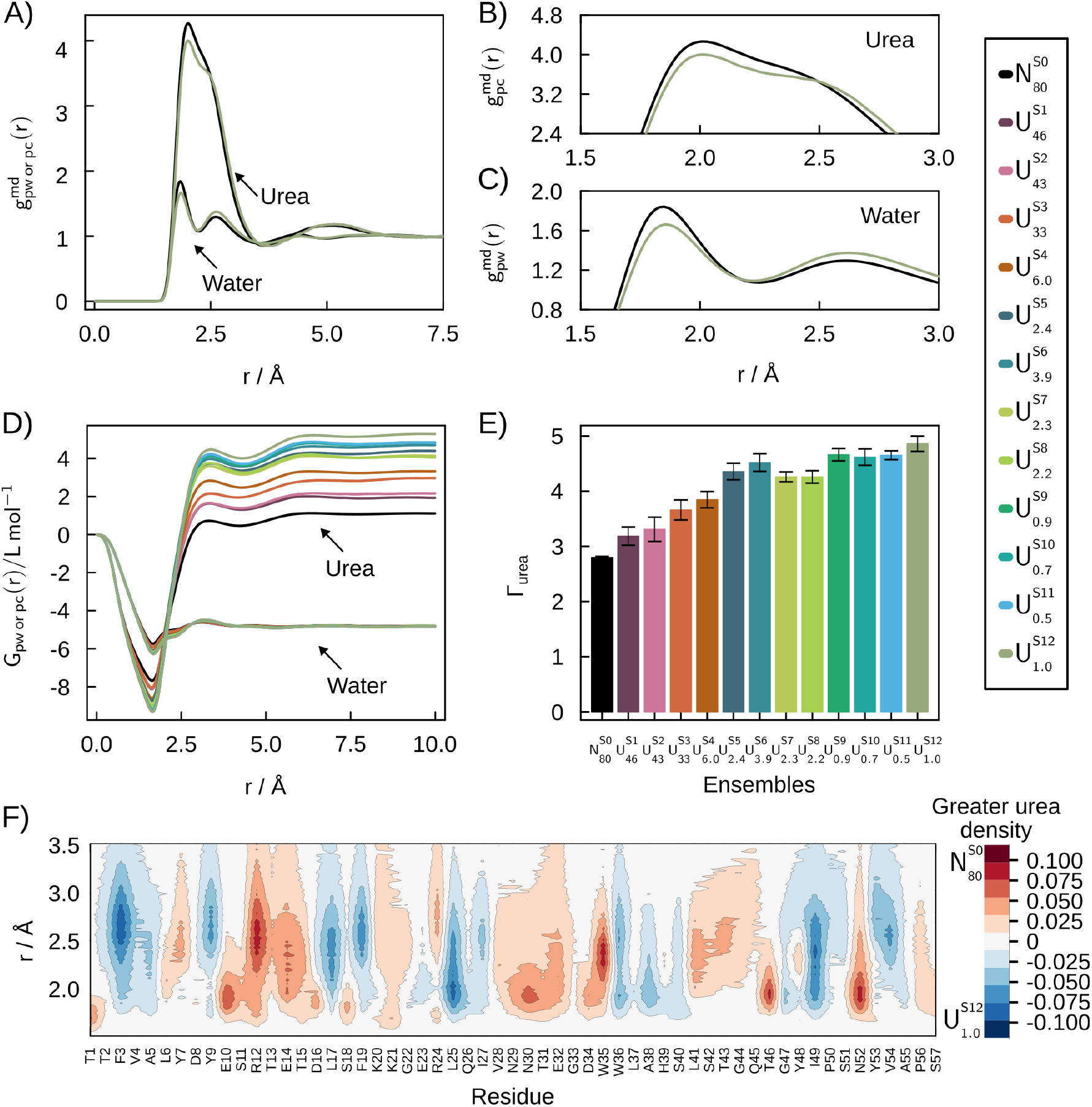
Solvation structures of the folding ensemble of the SH3 domain in 0.5 mol L^-1^ aqueous urea solutions. A) MDDFs of water and urea for the 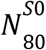 and 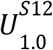 ensembles. B) and C) most prominent peaks in the MDDFs of urea and water. D) KBIs and E) Preferential interaction parameters (Γ). The preferential interaction of urea for unfolded states is greater than for the native 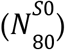 state and increases among denatured states. F) Difference in the breakdown of MDDF per residue in the vicinity of 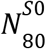 and the unfolded 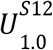 states. Red regions indicate greater urea density near the 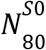 state, while blue regions show higher density around the 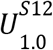 state.

The Kirkwood-Buff Integrals (KBIs) measure the overall solvent accumulation around the protein. The KBIs for urea, in Figure 2D, are greater than those of water, implying protein dehydration, as expected,^50–55^ and preferential interaction parameters with urea are always positive (Figure 2E). The distance-dependence of the KBIs, furthermore, provides insights on the contributions of interactions at each solvent shell to the build up of the final KBIs: The initial drop (Figure 2D) is associated with the excluded protein-urea volume, which is greater for denatured states. The urea KBIs then increase sharply, outgrowing those of water, and assume roughly the final converged value at 5 Å. Therefore, the accumulation of urea at the distances of H-bonds and nonspecific interactions are determining features leading to protein dehydration. Quantitatively, interactions above ∼2 Å have a greater contribution to final urea KBIs than interactions at H-bonding distances (for example, for the native state, from the minimum up to ∼2 Å the KBI increases ∼2.5 L mol^-1^, and from ∼2 Å to the final KBI it increased further ∼6 L mol^-1^, a difference that increases in denatured states). Evidence supporting these strengthened nonspecific interactions with unfolded states have been obtained for urea^14^ and other denaturants.^56^

The preferential interaction parameters (Γ_urea_) roughly increase monotonically with increasing protein denaturation, being correlated with the average fraction of native contacts (Q), β-sheet content, and solvent accessible surface area (SASA) (Supplementary Figure S6). For instance, 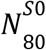 (the native state) has the lower Γ value compared with the other ensembles: it is 1.74 times lower than the higher Γ observed in The 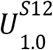 unfolded state. Thus, for SH3, the present results indicate that the variation of the chemical nature of the exposed surface upon denaturation does not need to be taken into account to qualitatively explain the urea denaturing effect.

An in-depth molecular picture of urea accumulation at the SH3 surface in different fold states can be obtained by a differential MDDF density map (Figure 2F). This map displays the difference in the contributions of each residue to the distribution functions of urea of the 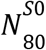 and 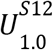 states. Red indicates a greater urea density in the native state, while blue indicates greater density in the unfolded state. Urea molecules are more densely found in the first shell of the 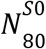 state around polar residues (threonines T1, T31, and T46, serines S18 and S57, and asparagines N30 and N52), charged residues (glutamate E10, arginine R12, lysine K21, and aspartate D16 and D34), and N- and C-terminal groups. Conversely, urea is found with greater density in the 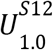 unfolded state at larger distances, around the hydrophobic residues such as phenylalanine (F3 and F19), tyrosine (Y9), leucine (L17 and L25), tryptophan (W36), isoleucine (I27 and I49), and valine (V54). Similar behavior is observed for water, indicating a competition between water and urea for specific and nonspecific interactions in the folded and unfolded structures (Supplementary Figure S8). Since urea was shown to have greater affinity than water to all but charged side chains,^21^ these observations reinforce the cooperation of urea binding and the exposure of the protein core along the denaturation mechanisms.

Red indicates greater TMAO density near the 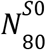 state, while blue indicates higher density around the 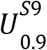 state.

Figure 3 shows the analysis of solvation structures of SH3 in aqueous solutions of TMAO. TMAO is a stabilizing osmolyte, a property associated with preferential exclusion (or, equivalently, preferential hydration).^2,57^ In Figures 3A and 3B we compare the MDDFs of the native state (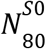 -black) with the unfolded state with greater TMAO exclusion (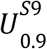 -green), and observe that the local density of TMAO in the denatured state is lower at H-bonding distances, but higher at distances associated to nonspecific interactions. Both TMAO and water, nevertheless, form specific interactions with the protein^24^ (H-bonds at ∼1.75 Å), but the TMAO peaks are much smaller than those of urea at these distances (Figures 2A and 2B). Also, there is a significant dip centered in ∼3.5 Å in the MDDFs of TMAO. This behavior is observed in all unfolded ensembles (Supplementary Figure S9). H-bonds with the protein involve the oxygen atom of TMAO (Supplementary Figures S10A and S10C). The second shell of the MDDF of the TMAO arises from its amphiphilic nature: TMAO has three methyl groups, which interact favorably with the hydrophobic regions of unfolded ensembles (Supplementary Figure S10C).

**Figure 3.**
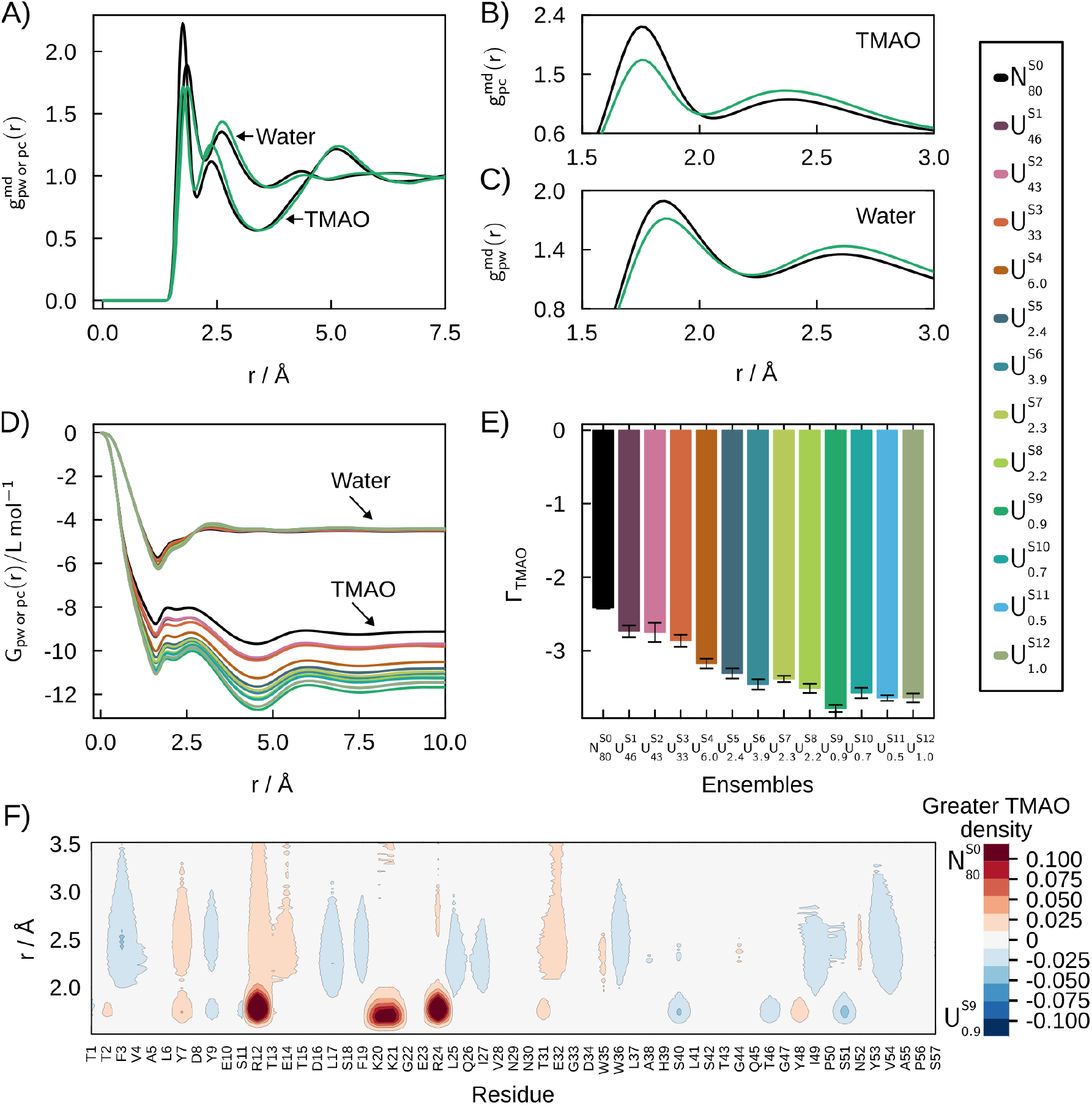
Solvation structures of the folding ensemble of the SH3 domain in 0.5 mol L^-1^ aqueous solutions of TMAO. A) MDDFs of water and TMAO for 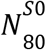 and 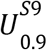 ensembles. B) and C) most prominent peaks in the MDDFs of TMAO and water. D) KBIs and E) Preferential interaction parameters. The exclusion of TMAO for unfolded states is greater than for the native 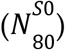 state and increases among denatured states. F) Difference in the MDDF of the urea in the vicinity of 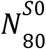 and of the unfolded 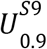 state.

Figure 3D displays the KBIs of water and TMAO for all SH3 states, as a function of the distance. Water KBIs are similar for all ensembles, but TMAO KBIs decrease with the increasing unfolding of the protein. TMAO exclusion at the short (< 2.0Å) and mid-range (∼2-5 Å) distances essentially determines the converged values of the KBIs. The smaller KBIs of TMAO relative to water imply that all the ensembles exhibit preferential hydration (Γ_TMAO_ < 0) (Figure 3E), and TMAO exclusion increases with denaturation. Once again, for SH3, the detailed nature of the exposed surface is not necessary to qualitatively infer the protecting action of the cosolvent.^2,57^ On the other hand, the present analysis shows that the more unfolded ensembles (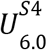 to 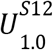) display greater TMAO exclusion, independently of the more hydrophobic character of the exposed protein core, which could lead to relatively greater TMAO affinity to the surface. Thus, perhaps counterintuitively, TMAO destabilization of denatured states is also cooperative.

Figure 3F shows the difference in the MDDF density per residue of TMAO in the vicinity of 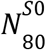 and 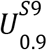. Red and blue indicate higher TMAO density around 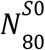 and 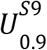 respectively. The density of TMAO around 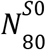 and unfolded 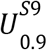 are similar for most residues. Interactions in the first solvation shells of R12, R24, K20, and K21 are responsible for the greater MDDF peak at ∼1.75 Å in the native state (Figure 3B). The increase in density of the second solvation shell in the 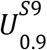 is primarily linked with nonpolar (F3, L17, L25, W36, I49, and V54) residues. Differential water density maps resemble those observed in the urea solutions (Supplementary Figures S8 and S11). Thus, water accumulates around the same residues independently of the cosolvent, albeit with a higher density in the TMAO solutions.

The MDDFs can be decomposed into the contributions of the backbone and side chains of the protein, as shown in Figure 4. The H-bonds responsible for the first peak of the urea MDDF have important contributions from both backbone and side chains, while for TMAO the backbone contribution is negligible. Notably, the peak associated with urea-backbone interactions becomes slightly greater for the denatured state, such that these bonds are relatively more frequent in the denatured states relative to other interactions. Interactions with the side chains are more complex: at short distances, the contributions of side chains to urea binding decreases (dashed vs solid green lines in Figure 4A), but they slightly increase for nonspecific interactions, mostly with nonpolar residues (Supplementary Figure S7). The increased dehydration promoted by urea is then a combination of its greater association with the backbone at short distances, and with nonpolar residues exposed upon denaturation. TMAO preferential exclusion, on the other hand, is mostly driven by similar interaction patterns in native and denatured states, with quantitative accumulation driven by the surface area exposed in each state.

**Figure 4.**
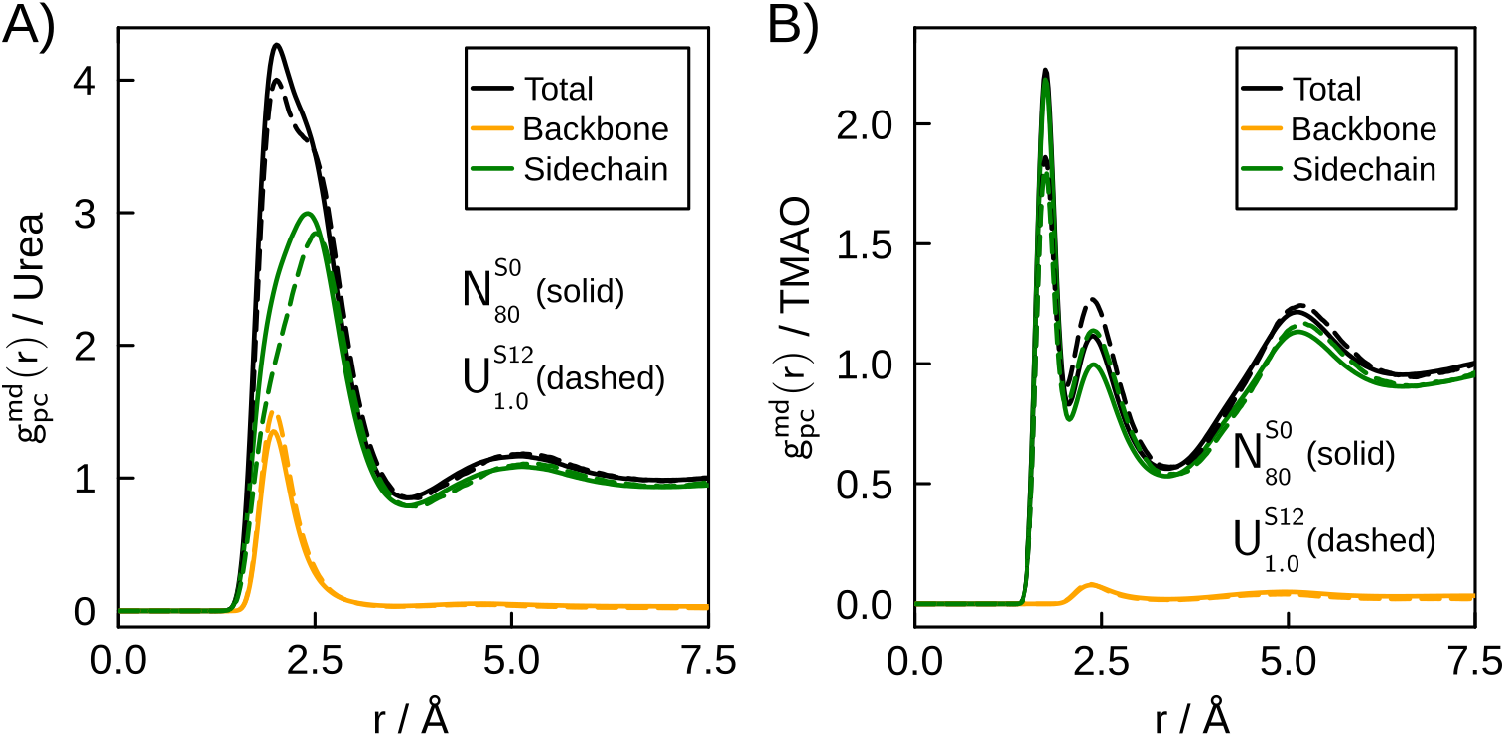
Decomposition of the MDDFs of urea and TMAO relative to SH3 in backbone and side chain contributions for the native (solid line) and a highly denatured state (dashed line).

### Osmolyte effects on the folded and unfolded states of BdpA

In this section we will discuss the solvation structure of BdpA, a model α-helix protein, in the presence of urea and TMAO. On the one hand, the overall solvation properties of BdpA are similar to those of SH3: in Figure 5A and 5B, density augmentations of water and urea at short distances from the protein can be observed. Figure 5B shows that the H-bonding peaks are greater in the native state relative to the denatured state, but nonspecific interactions are strengthened in the denatured states. Qualitatively similar MDDF profiles are observed for TMAO in Figures 6A and 6B. Urea displays greater KBIs than water (Figure 5D) and solvates preferentially the BdpA domain (Figure 5E). TMAO, on the contrary, is preferentially excluded (Figures 6D and 6E). Urea is a better solvent for SH3 than for BdpA, and TMAO is a poorer one, as judged by the preferential interaction parameters. This is consistent with the observation that urea predominantly unfolds β-sheet structures.^58–60^

**Figure 5.**
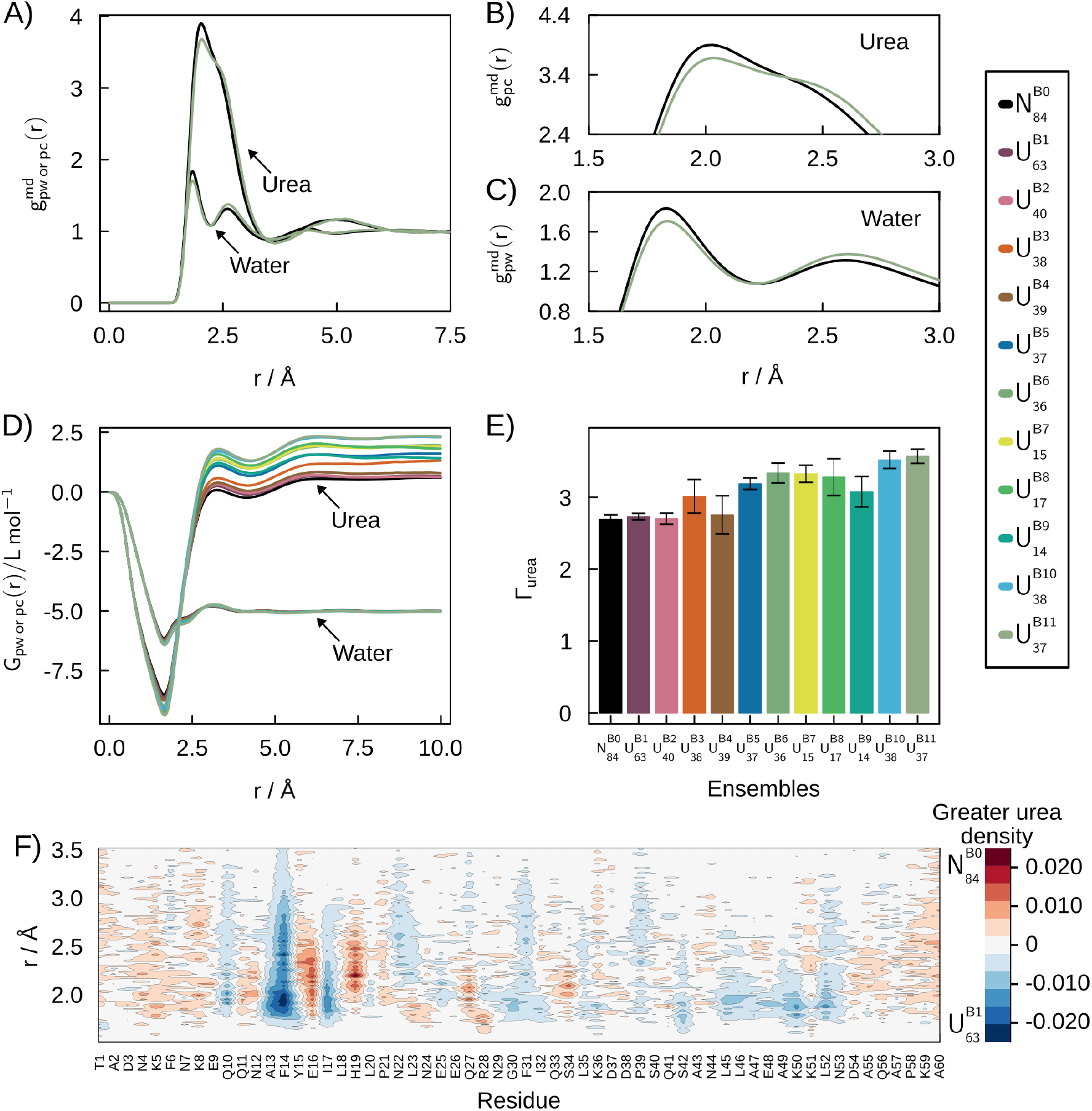
Solvation structures of the folding ensemble of BdpA in 0.5 mol L^-1^ aqueous urea solutions. A) MDDFs of water and urea for 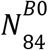 and 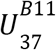 ensembles. B) and C) most prominent peaks in the MDDFs of urea and water. D) KBIs and E) Preferential interaction parameters. The preferential interaction of urea for unfolded states is greater than for the native 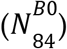 state and increases among denatured states. F) Differential density map of the MDDF breakdown per residue of 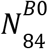 and 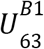. Red indicates greater urea density around 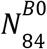 and blue around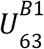.

**Figure 6.**
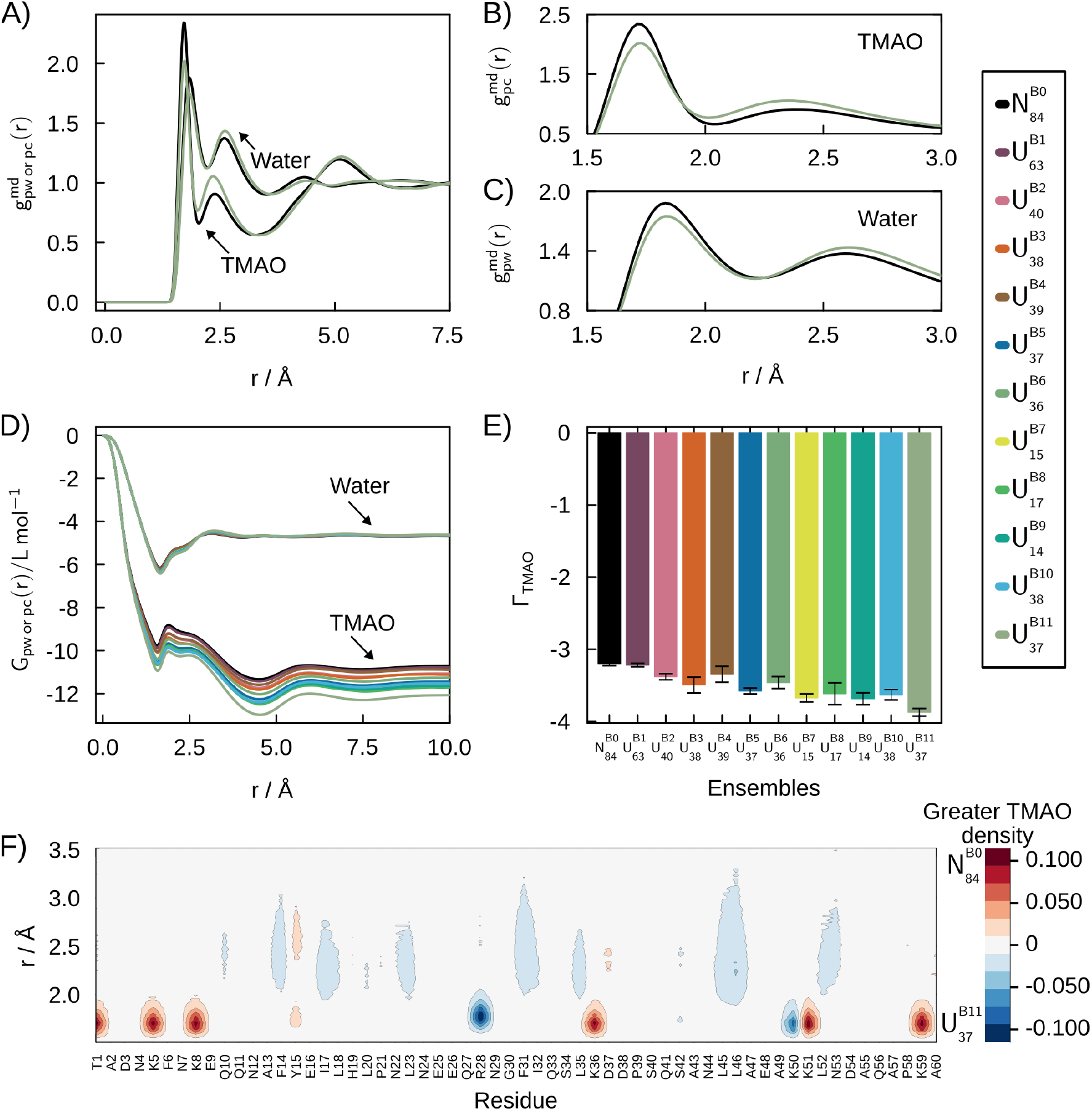
Solvation structures of BdpA in 0.5 mol L^-1^ aqueous solutions of TMAO. A) MDDFs of water and urea for 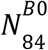 and 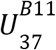 ensembles. B) and C) most prominent peaks in the MDDF of TMAO and water. D) KBIs and E) Preferential interaction parameters. The preferential interaction of TMAO for unfolded states is lower than for the native 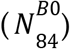 state and decreases for denatured states. F) Differential density map of the MDDF breakdown per residue of 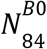 and 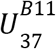. Red indicates greater TMAO density around 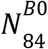 and blue around 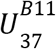.

On the other hand, what is notable about BdpA is that its first four denatured subsets, 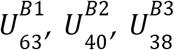, and 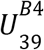, display preferential interactions roughly equivalent to those of the native 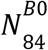 state, notably in urea (Figure 5E) but also in TMAO (Figure 6E). The 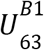 state is, effectively, only partially denatured: it displays 63% of the crystallographic helical content and 72% of native contacts, versus 84% and 77% of the 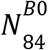 ensemble from the folded basin (Supplementary Table S1). The partial denaturation of 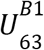 is associated then to its secondary-structure compositions (Figure 1F), but not so to its surface area. 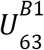 is an important denaturation intermediate of BdpA: it is characterized by the partial denaturation of helix I (Supplementary Figure S5). Helix I is known to be labile in BdpA, providing structural variability to the folded ensemble and leading the initial stages of denaturation.^42,45,46^ By contrast, the 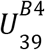 set, with 39% of helical content and 54% of native contacts, is much further denatured. Yet, its surface accessible surface area, 52.7 nm^2^ (Supplementary Table S1) is also similar to that of the native set, of 51.2 nm^2^, indicating that it has conserved a globular fold despite the rearrangement of the secondary structure and tertiary contacts. These first four BdpA denatured sets have SASAs within 4% of that of the native ensemble, while the remaining unfolded ensembles (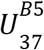 to 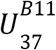) have accessible surface areas that are at least 11% greater than that of 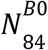. Therefore, while the SASA values determine the resulting preferential interaction parameters with urea, the denatured ensemble of BdpA cannot be reduced to this parameter, and this will have important consequences to transfer free energies.

Figure 6F shows the differential density map of 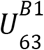 and 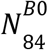 per residue. The exposure of F14 and neighboring residue to urea highlights the link between helix I denaturation and the solvation structure. The pronounced exposure of F14 to urea is a common feature of all denatured ensembles, except of the unfolded but compact 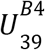 (Supplementary Figure S12), where it remains buried in the protein’s hydrophobic core. Variations in the density of TMAO around each residue are smaller due to its preferential exclusion. However, in denatured states, TMAO interacts more with hydrophobic residues, and less with the charged residues, particularly lysines (Figure 6F and Supplementary Figure S13).

In summary, BdpA interactions with urea and TMAO are qualitatively similar to those of SH3, but two sets of ensembles can be distinguished: i) partially denatured states with a slight increase in surface area interact with the solvent similarly to the native state; ii) denatured states with greater surface area are dehydrated by urea and preferentially hydrated in the presence of TMAO. These contrasting behaviors contribute to a multifaceted solvent dependency of the denaturation equilibrium of BdpA, which will be quantitatively discussed below.

### Osmolyte induced shifts in denaturation equilibrium constants

The variation of the equilibrium constant, *K*, of among two folding states *U* and *N*, as a function of the osmolyte activity *a*_*c*_, can be obtained with the Wyman linkage equation,^49,61^

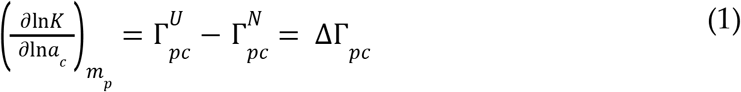

where Γ are the preferential interaction parameters. ΔΓ_*pc*_ greater than zero implies that an increase in cosolvent concentration favors the denatured state. In qualitative terms, if the cosolvent solvates more favorably the denatured state than the native state, increasing the cosolvent concentration will shift the equilibrium towards denaturation.

Figure 7 displays ΔΓ for SH3 and BdpA in urea and TMAO at 0.5 mol L^-1^, considering the equilibrium involving the native states and each of the unfolded states. The denatured states of SH3 are always stabilized by urea addition (Figure 7A - ΔΓ positive) and destabilized by TMAO (Figure 7B). For BdpA, ΔΓ is also always positive for urea, although within the modeling uncertainties for the first four partially unfolded states. Finally, TMAO favors the native state of BdpA (ΔΓ negative) relative to all unfolded states, with perhaps the exception of 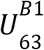. In all cases, the osmolyte effects are more prominent as the degree of denaturation increases.

**Figure 7.**
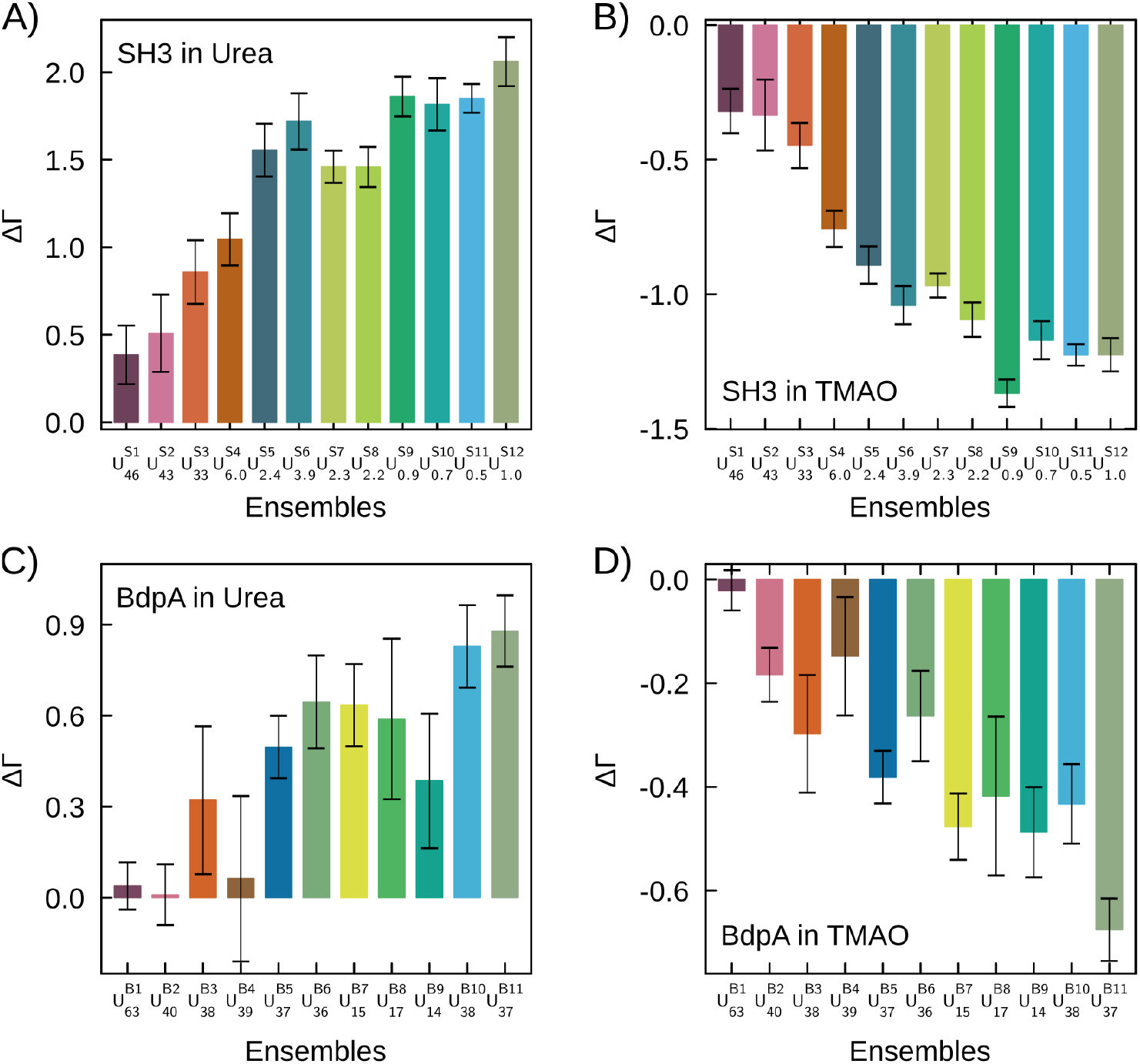
ΔΓ for each folding ensemble of structures of SH3 and BdpA proteins, relative to the native-state clusters. Panels A) and B) show ΔΓ values for each ensemble of SH3 in urea and TMAO 0.5 mol L^-1^. Similarly, panels C) and D) show ΔΓ values for each ensemble of BdpA under the same cosolvent conditions.

Thus, in general, increasing the concentration of urea favors denatured states, while the native state is favored by TMAO, consistently with the overall known roles of these osmolytes on protein stability.^62^ Nevertheless, the rule breaks down for denatured states displaying similar solvent accessible surface area than the native state. The linkage relation, Equation 1, does not allow, however, obtaining the absolute dependence of the equilibrium constant of denaturation (or, equivalently, on the denaturation free energy) as a function of the cosolvent concentration. It is necessary to integrate the concentration-dependence of the free energy for each state from pure water to the final ternary solution to obtain the net effect of the cosolvent on the folding equilibrium. This computation was performed for selected states, as described below.

### Osmolyte dependence of unfolding free energies for selected denatured states

The absolute effect of a cosolvent on the protein folding equilibrium can be determined by the difference of equilibrium constants of folding at each solvent concentration, relative to water. This corresponds to the variation of the free energy of folding upon cosolvent addition. Experimentally, these were measured by NMR with various osmolytes, including urea and TMAO,^15^ for the SH3 domain from the *Drosophila signal transduction protein* (Drk-SH3), which shares the same fold of the SH3 model here studied.^63^ Alternatively, if the folding states can be stabilized in the time-scale of the experiment, transfer free energies of each state from water to the cosolvent solution can be obtained, from which the same concentration dependence of the folding free energy can be obtained.^2^ These two approaches correspond to the reaction pairs of the Tanford Transfer thermodynamic cycle,^64,65^ shown in Figure 8.

**Figure 8.**
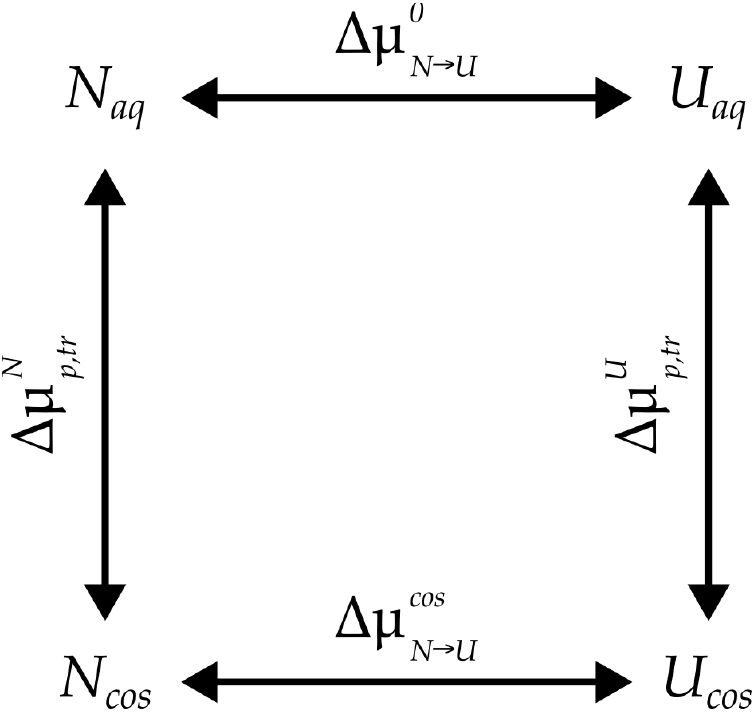
Thermodynamic cycle of the Tanford Transfer Model.^64,65^ This cycle illustrates how the folding free energies between the native (N) and unfolded (U) states in the presence of a cosolvent (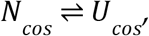 given by 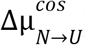) differs from that in water (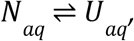 given by 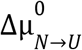). Direct measures of equilibrium constants with and without cosolvent (horizontal reactions) allow the computation of the cosolvent effect on the folding free energy, 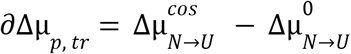. Alternatively, the transfer free energies for the native 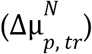 and unfolded 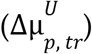 states can be obtained (vertical reactions), and 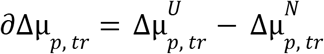.

Since computational modeling of the folding equilibrium of even small proteins such as SH3 and BdpA is difficult, the effect of the cosolvents on the folding equilibrium will be estimated by computing transfer free energies.^1,2,10,66^ The transfer free energy,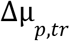 of a structure from pure water to a solution of the cosolvent is

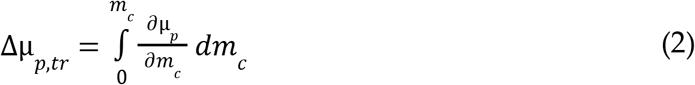

Where 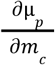 is the variation of the chemical potential of the structure with cosolvent concentration, and the integral is performed from pure water to the desired *m*_*c*_ concentration of the cosolvent. This computation mimics the experiment of adding a cosolvent to an aqueous solution of a protein while roughly preserving the protein folding state, either folded or an unfolded construct.^2^

Transfer free energies can be obtained by numerical integration of Equation 2, with the integrand computed from preferential interaction parameters and activities obtained at various cosolvents concentrations using,^2^

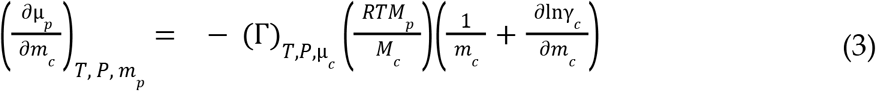

where γ_*c*_ is the cosolvent activity, and *M*_*p*_ and *M*_*c*_ are the molar masses of the protein and the cosolvent, respectively. The activities of urea and TMAO can be obtained from experimental data,^2^ and the preferential interaction parameters can be obtained by simulating a protein model in different cosolvent concentrations.

We focus on three folding states for each protein: the native states (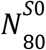 for SH3 and 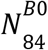 for BdpA) and two unfolded states. The unfolded states were chosen from the ΔΓ extremes in urea or TMAO at 0.5 mol L^-1^ (Figure 7). For SH3, we selected 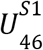 and 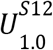, while for BdpA, 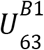 and 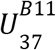 were selected (see Figures 1 and 2). Simulations were performed for the most representative structure of each ensemble in 7 concentrations of urea and TMAO ranging from 0.1 to 1 mol L^-1^. Solution structures were characterized using MDDFs and the KB theory, as previously described, to obtain 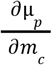 using Equation 3. Preferential interactions obtained are shown in Supplementary Figure S16. The transfer free energy of each model (Δμ_*p, tr*_) was then obtained from the numerical integration of Equation 2. Finally, the effect of the cosolvent in the folding free energies are 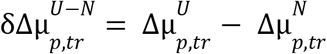, the difference in transfer free energies associated with the N ⇌ U transition, for each unfolded state.

Figure 9A shows that the variation in unfolding free energy of SH3 states is negative for all urea concentrations, and thus the denatured states are favored. 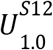 is a highly unfolded state, with a large surface area, and consistently becomes stabilized by urea to a greater extent than the more compact 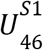 model. In the presence of TMAO (Figure 9B), a positive ∂Δμ_*p, tr*_ implies destabilization SH3 unfolded states, consistently with the “osmophobic effect”.^8^ ∂Δμ_*p, tr*_ for 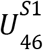 is smaller than for 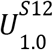 because TMAO favors compact structures.

**Figure 9.**
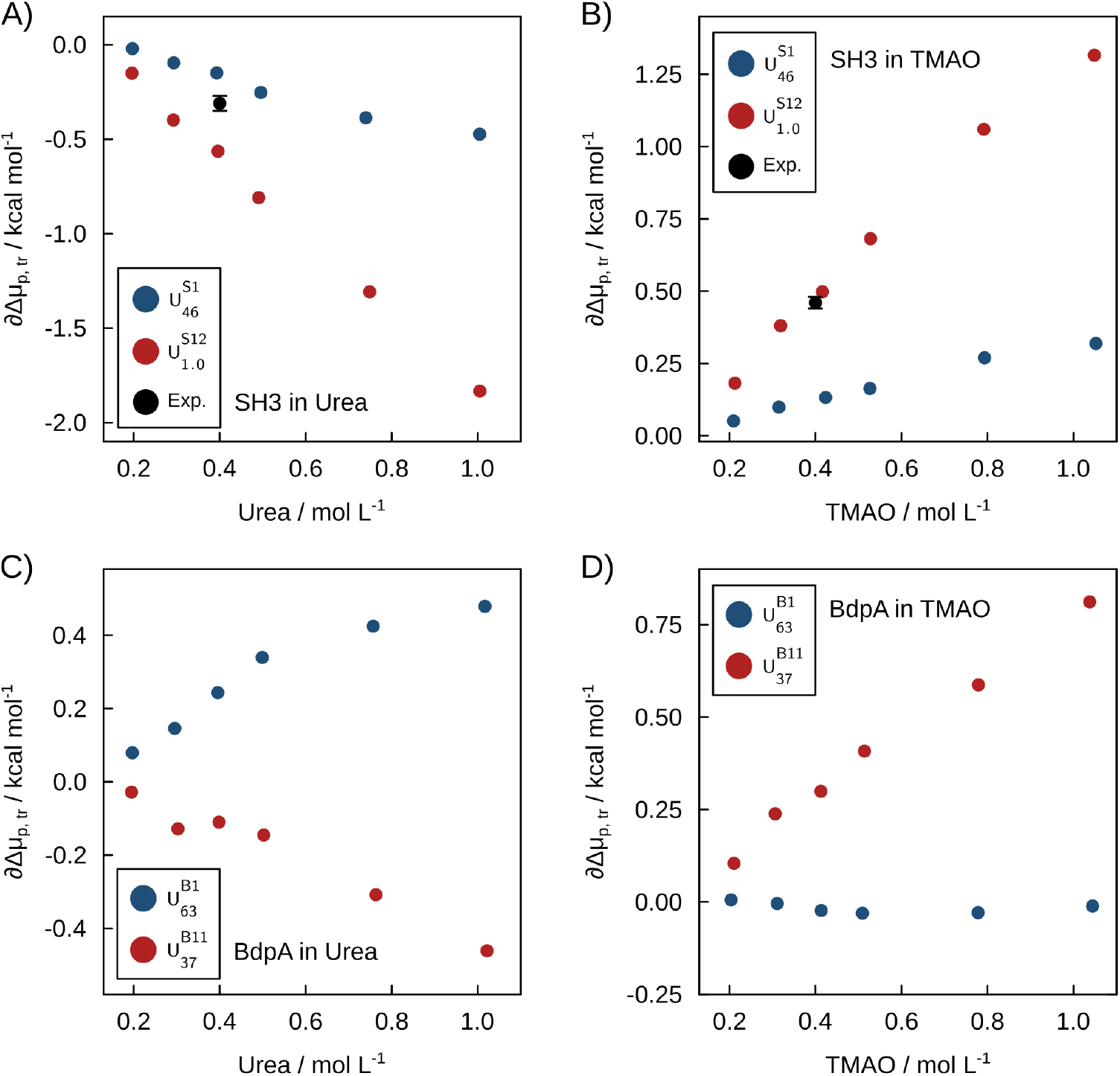
Variation of folding free energies (∂Δμ_*p, tr*_) relative to pure water as a function of cosolvent concentration, for selected denatured states. A) and B) ∂Δμ_*p, tr*_ for SH3 models in urea and TMAO, respectively. This analysis focused on the 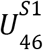 (blue) and 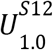 (red) unfolded ensembles. C) and D) ∂Δμ_*p, tr*_ for BdpA for the folding equilibrium associated to the 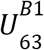 (blue) and 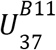 (red) BdpA ensembles. Black symbols indicate experimental data for Drk-SH3.^15^

Experimentally, the free energy of Drk-SH3 folding equilibrium was shown to be affected by urea, at 0.4 mol L^-1^, by -0.31 ± 0.02 kcal mol^-1^.^15^ At this concentration, the computed variations of unfolding free energies of SH3 upon urea addition are -0.15 and -0.56 kcal mol^-1^ for the 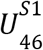 and 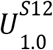 states, respectively. Similarly, the experimentally determined variation in folding free energy in 0.4 mol L^-1^ TMAO solutions was 0.46 ± 0.02 kcal mol^-1^, and here we obtain 0.13 and 0.50 kcal mol^-1^. Experimental values are, thus, within the lower and upper bounds obtained for the effect of the cosolvents on the folding equilibria. This qualitative agreement in the sub-kcal mol^-1^ energy regime is a remarkable validation of the folding ensemble obtained by the simulations, the force-fields used for representing solvent-protein interactions, and the methods used for solvent structure analysis. The experimental 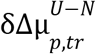 are shown as black dots in Figure 9.

A similar analysis of cosolvent effects on the folding equilibria of BdpA ensembles 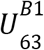 and 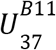 is shown in Figures 9C and 9D. 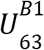 has positive 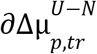 in urea and virtually zero in the presence of TMAO. For the 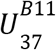 state, with greater denaturation and surface area, folding free energies confirm the denaturant and protector effects of urea and TMAO (Figures 9C and 9D - red dots). In the previous section, we showed that the solvation characteristics of the 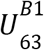 set are similar to those of the native state of BdpA. Here, we show that this is a particular case where urea exerts a destabilizing effect on a partially unfolded state, while in TMAO the unfolding equilibrium involving this state is essentially independent of the cosolvent concentration. This observation was only partially deducible from the Wyman linkage relations in Figures 7C and 7D, considering the standard error of ΔΓ values. We cannot obtain the experimental population ratio of 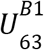 -like states vs native states from our simulations, but in our SBM model 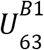 was twice as populated as 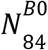. This implies a free energy difference of roughly -0.4 kcal mol^-1^ at 300K. The urea effect on the folding equilibrium, computed here, is in the same order of magnitude, but favoring the native state. Thus, it could significantly reduce the population of the partially denatured helix I, redirecting unfolding along new pathways.

Transfer free energies can also be estimated by semi-empirical models, for example using the assumption of backbone and side chain additivity^67,68^ and effective estimates of the solvent exposure of residues in random-coil models of denatured states.^68–71^ From these models Bolen and co-workers deduced that osmolyte-backbone interactions are dominant for the protecting or denaturing effects.^3–6,72,73^ We can’t decompose the transfer free energies obtained here in group contributions, but the analysis of the distribution functions points to favorable urea-backbone interactions and the TMAO-backbone exclusion, while, at the same time, suggesting that nonspecific interactions also play a crucial role in the final cosolvent preferential interactions.

## Discussion

In this work, we developed a computational approach to characterize the effect of osmolytes on protein folding ensembles. Our pipeline combines SBMs and all-atom simulations to model the folding landscapes of SH3 and BdpA, and the solute-solvent interactions of every conformation. To elucidate the solvation structures and quantify the stabilization promoted by urea and TMAO, we employed MDDFs, which are crucial for the characterization of solvation of complex molecular shapes. We then applied the Kirkwood-Buff theory to obtain preferential interactions and the osmolyte dependence of folding free energies.

Our results confirm urea’s preferential interaction and TMAO’s preferential exclusion from protein surfaces, while adding a new dimension to how these interactions vary across the protein folding landscape. Different denatured states exhibit distinct interactions with water and cosolvents, despite significant correlations with solvent accessible surface area. Globular unfolded states can be only marginally affected by osmolytes, with urea potentially promoting their destabilization. MDDFs reveal that water, urea, and TMAO form both H-bonds and nonspecific interactions with the protein surface (Figures 2A-C and 3A-C). Specific interactions are stronger in the native states, whereas the nonspecific interactions are strengthened in the denatured states. These patterns hold across different solvent components, protein types, and folds, reflecting the increased exposure of hydrophobic core residues upon denaturation.

Solvent exposure, fraction of native contacts, and the secondary structure elements (β-sheet and α-helix) are key determinants of preferential interactions. In the case of SH3, for which these properties are highly correlated, preferential interactions and folding free energies vary monotonically with increasing denaturation. The qualitative understanding of the unfolding paths does not depend then on the exact nature of the denatured states. On the contrary, in the case of BdpA, the secondary structure content does not directly correlate with the solvent accessible surface area or the fraction of native contacts. Thus, denatured states like 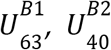, and 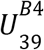 exhibit solvation properties similar to the native state, despite their lower helical content.

Simulating representative structures of selected folding states in various TMAO and urea concentrations allowed us to compute the absolute osmolyte effect on the folding free energies. Notably, the experimentally determined folding free energy dependence on urea and TMAO for an SH3 homologous protein is found within the computed bounds, validating the force-fields, modeling strategy, and solvation analysis methods. At the same time, the solvation of a single denatured state is not sufficient to infer quantitatively the role of cosolvents in the folding equilibrium, consistently with the unfolded ensemble variability playing a key role.^74^

## Conclusions

Understanding the molecular mechanisms by which osmolytes modulate protein stability across the entire folding landscape, including the challenging-to-characterize denatured states, remains a critical goal in biophysics. This work addressed this challenge by employing a novel computational pipeline combining coarse-grained and all-atom simulations with Minimum-Distance Distribution Functions (MDDFs) and Kirkwood-Buff theory to dissect the thermodynamic effects of urea and TMAO on the β-sheet SH3 and α-helical BdpA domains.

We demonstrated that urea generally stabilizes unfolded states through preferential dehydration driven by strong hydrogen bonds with the protein backbones and favorable nonspecific interactions, while TMAO enhances preferential hydration, consistent with its protective effect. Crucially, these effects are conformation-dependent: urea significantly destabilizes specific partially unfolded intermediates of BdpA that remain largely unaffected by TMAO, highlighting the potential for osmolytes to selectively influence folding pathways. This detailed view was enabled by the first calculation, to our knowledge, of transfer free energies for a wide range of denatured states directly from atomistic simulations, yielding folding free energies consistent with experimental data and validating our approach.

While our study focused on two specific proteins and two osmolytes, the results underscore the necessity of moving beyond native-state analyses to capture the complex interplay between solutes, cosolvents, and proteins across the conformational ensemble. The insights gained emphasize the role of non-polar, dispersive forces in modulating solvation thermodynamics. Future work could extend this methodology to explore a broader range of osmolytes and mixtures, investigate different protein architectures, assess the influence of force field parameters, and further dissect the specific contributions of residue types to these interactions. Ultimately, this work provides a more nuanced understanding of osmolyte action and offers a robust framework for investigating macromolecular solvation in complex environments, contributing to efforts in protein engineering and solvent design.

## Methods

### Structure-Based Models (SBMs) simulations and analyses

Folding ensembles of the SH3 domain (PDB: 1FMK)^28^ and BdpA (PDB: 1BDD)^29^ were generated from Cα-Structure-Based Models simulations.^75,76^ The contact map of the native structures was determined using the *Contact of Structural Units* (CSU) algorithm,^77^ resulting in 140 and 98 contacts, respectively. The Cα-SBMs were generated using the SMOG web server (https://smog-server.org/)^78^ and simulations were performed with Gromacs 4.6.7.^79^

The folding ensemble described for BdpA in our previous publication was used in this work.^42^ A similar protocol was adopted here to obtain the SH3 folding ensemble. For the SH3 protein, the simulations were conducted in temperatures ranging from 110 to 170 units, with a temperature step of 10 units. Simulations at constant temperatures consisted of 5 x 10^8^ steps with a time step of 0.0005 reduced units. Once the temperature of maximum specific heat, C_v_, was roughly identified, a new set of simulations with temperatures varying between 135 and 139 with 1 unit temperature steps was performed, to localize within ∼1 unit the folding temperature (T_f_). In reduced temperature units, the simulations were performed within 0.91 and 1.41 with 0.083 steps and within 1.12 and 1.15 with 0.0083 temperature unit steps for the first and second sets of simulations. The simulation at the T_f_ was extended ten times to increase the sampling of conformational transitions.

The temperature dependence of the specific heat, C_v_(T), and the potential of mean force as a function of the fraction of native contacts, F(Q), of these simulations were obtained with WHAM,^80^ as implemented in SMOG2.^81^ A contact was classified as native if the distance between the corresponding Cα atoms did not exceed 20% of the value observed in the experimentally determined models. The statistical convergence of the Q-values at the T_f_ was confirmed by block-averaging using the MolSimToolkit.jl package (Supplementary Figure S2).

### Protein folding phase-space

The *Energy Landscape Visualization Method* (ELViM)^35,44,82–86^ was used to represent the folding phase space of SH3 and BdpA. The ELViM method uses matrices of internal distances of conformations to establish a reliable structural similarity metric without predefined reaction coordinates. The similarity matrix is projected into a 2D space optimizing the distances between points to be correlated with the dissimilarity between structures. For this projection, we apply the Force Scheme technique,^87^ as implemented in ELViM.^35^ The secondary structures were computed with the DSSP method,^88,89^ using the ProteinSecondaryStructures.jl package.

### Atomistic simulations and analysis

All-atom representations of 5,000 SBM models for each protein, extracted from the simulations performed at the T_f_ were reconstructed using the Pulchra software.^90^ We have studied the solvation of each reconstructed structure in solutions of urea 0.5 mol L^-1^ and TMAO 0.5 mol L^-1^.

Orthorhombic simulation boxes were constructed with a minimum distance of 12.0 Å from the protein extrema using Packmol.^91,92^ Details of each system, including the box volume and the number of solvent molecules, are shown in Supplementary Tables S2-S5. Harmonic potentials with 25 kcal mol^-1^ force constants were applied to the Cα atoms to preserve the Cα-SBM topology during minimization, equilibration, and production, while allowing relaxation of the reconstructed atoms and solvent. The CHARMM36 force field^93^ for the protein and the TIP3P water model^94^ were used. Urea molecules were modeled with the combination of CGenFF^95,96^ and the charges of the KBFF force-field, which was developed to reproduce the KBIs of urea and water molecules.^16^ TMAO was described using the Netz force field^97^ with the scaling correction proposed by Shea and co-workers,^98^ designed to accurately reproduce the Kirkwood-Buff integrals (KBI) of TMAO in ternary solutions containing water, TMAO, and ribonuclease T1 (RNase T1).^2^

All atomistic simulations were performed in Gromacs 2021.2^99^ at 298.15 K, 1 atm, and with a time step of 2 fs. Initially, the systems were minimized by up to 20,000 Steepest Descent steps and equilibrated for 1 ns in constant-volume and constant-temperature ensemble (NVT) followed by 1 ns of constant-volume and constant-pressure (NPT) simulation. Temperature and pressure were controlled using the modified Berendsen thermostat^100^ and Parrinello-Rahman barostat.^101^ Finally, 10 ns production simulations were performed in the NPT ensemble for each system, totaling 240 µs of simulation, used to obtain MDDFs, KBIs and preferential interaction parameters of Figures 2 to 6, and the data of Figure 7.

Simulations in urea and TMAO at 0.1, 0.2, 0.3, 0.4, 0.5, 0.75, and 1.0 mol L^-1^, of the 6 selected representative structures of BdpA and SH3 ensembles, were performed to determine the transfer free energies used to obtain Figure 9. (see the next section for more details). 20 replicas of 25 ns were performed for each system, totalling 42 µs of simulation (Supplementary Tables S4 and S5). Representative structures were selected by the *most_representative_structure* function in the MolSimToolkit.jl package version 1.23.0. The most representative structure has the minimum RMSD relative to the average structure of each set.

### Characterization of the solvent structure and thermodynamics

This study employs a notation system for clarity: w (water), p (protein), c (cosolvent), and s (water or cosolvent). We developed the ComplexMixtures.jl package^25^ to characterize the solvation structures from Minimum-Distance Distribution Functions (MDDFs)^24^ and the Kirkwood-Buff theory of solvation.^102^ Specifically, the package computes the distribution functions, Kirkwood-Buff Integrals (KBIs), and atomic, group, and residue decompositions required for a molecular understanding of solvation. Density maps like that of Figure 2F were generated with the *ResidueContributions* function of ComplexMixtures.jl. From the KBIs, preferential interaction parameters (Γ) and the Wyman-linkage relation (Equation 1) can be obtained.^49^ The computation of transfer free energies followed the protocol described by Lin and Timasheff.^2^ The combination of MDDFs, Γ, ΔΓ, and Δµ parameters comprehensively characterizes the solvation of the folding ensembles.

MDDFs are defined in terms of the average number density of solvent molecules *n*_*c*_(*r*) in the simulation, relative to the density of an ideal gas distribution 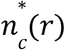,

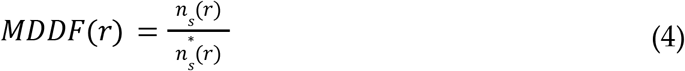

where *r* is the minimum-distance between the protein and the cosolvent. The ideal distribution of minimum-distances has to be generated numerically to account for the shape of solute and solvent. Further discussion about MDDFs and their importance in the study of solvent structures are found in your previous publications.^24–26,103^

With an adequate reference state, the minimum-distance counts can be used to compute the Kirkwood-Buff integrals (KBIs) and preferential interaction parameters (Γs). In a finite sub-volume of the system, the KBIs is given by Equation 5.

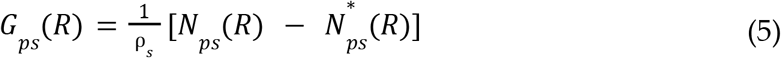

where *N*_*ps*_(*R*) and 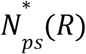 are the number of minimum distances between the protein and the solvent within *R*, and the number of equivalent distances within *R* in an ideal system. The KBIs are the limit of *G*_*ps*_ (*R*) for large *R*, and are the excess volume occupied by the cosolvent relative to the volume that it would occupy in the absence of solute-solvent interactions.

The thermodynamic property that measures if the cosolvent accumulates or is depleted from the protein domain, compared with the bulk solution, is the preferential interaction parameter (Γ). In the infinitely dilute solute limit, the cosolvent preferential binding to the protein (Γ_pc_) can be approximated by site counts in a finite volume defined by *R*, with

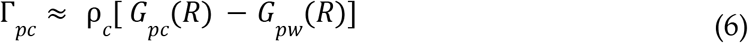

where *R* is large enough to encompass the portion of the solution affected by the presence of the solute.

Given Γ_pc_ for two states, the Wyman linkage equation^49^ (Equation 1) is used to obtain the direction into which a cosolvent will drive the reaction at a given cosolvent concentration. Transfer free energies are computed using the numerical integration of Equation 2, using Equation 3 as the integrand, from a finite set of simulations of the states considered in different concentrations of the cosolvent. Notebooks with reproducible calculations of the transfer free energies shown in Figure 9 are available at https://m3g.github.io/PereiraMartinez2025.jl. The average of preferential interaction parameters from 20 simulations, calculated across seven cosolvent concentrations, are used to determine ∂Δμ_*p, tr*′_ , and are summarized in Supplementary Figure S16. The Plots.jl^104^ package was used to create figures.

## Author information

### Authors and Affiliations

Institute of Chemistry and Center for Computing in Engineering & Science, Universidade Estadual de Campinas (UNICAMP), 13083-861 Campinas, SP, Brazil

Ander Francisco Pereira and Leandro Martínez

### Contributions

A. F. P. proposed strategies for, and executed, simulations and data analysis, and wrote the paper. L. M. conceived the project, supervised research, developed software, and wrote the paper.

## Supplementary Information

Structural characterization of folding ensembles (Table S1); SBM folds space characterization (Figures S1 and S2); Representative structure of each fold subset (Figure S3); Secondary structure content per residue in each subset (Figures S4 and S5). Correlation between Γ and SASA (Figure S6); MDDF decompositions of urea and TMAO by backbone and side chain contributions (Figure S7); Density maps for water (Figure S8 and S11); MDDFs for all ensembles for SH3 and TMAO (Figure S9) and group contributions (Figure S10); Residue contributions differential density maps for all states in urea and TMAO (Figures S12 to S15). Γ for selected folds at multiple concentrations (Figure S16); Simulation box details (Tables S2 to S5).

## Data Availability

Data and codes necessary for the computations here reported and detailed reproduction of the computation of transfer free energies are available at https://m3g.github.io/PereiraMartinez2025.jl.

## Acknowledgments

The authors acknowledge the financial support of FAPESP (2018/24293-0, 2013/08293-7, 2018/14274-9, 2019/17874-0, 2020/04549-0), and CAPES 206-04/092018. Research developed with the help of CENAPAD-SP (National Center for High-Performance Processing in São Paulo), project UNICAMP / FINEP - MCTI, and the Coaraci Supercomputer.

## Table of Contents

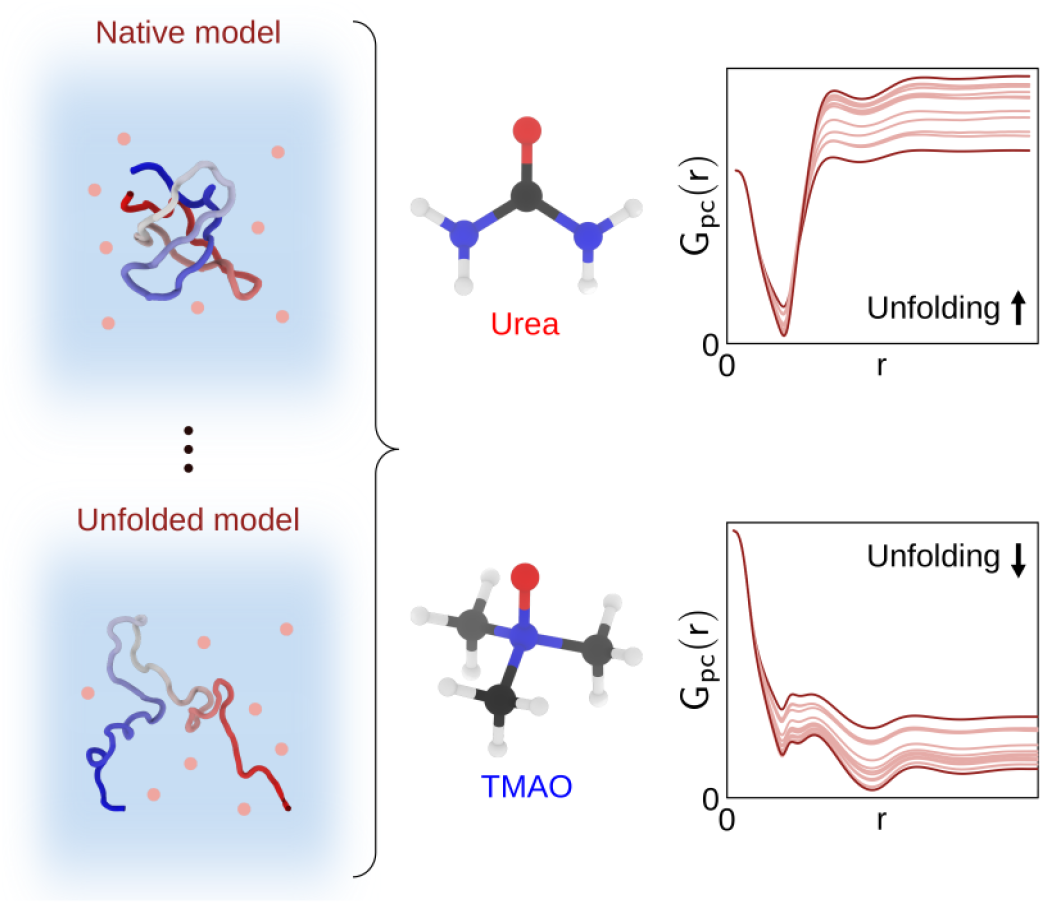

## References

(1) Bolen, D. W.; Rose, G. D. Structure and Energetics of the Hydrogen-Bonded Backbone in Protein Folding. Annu. Rev. Biochem. 2008, 77, 362–.

(2) Lin, T. Y.; Timasheff, S. N. Why Do Some Organisms Use a Urea-Methylamine Mixture as Osmolyte? Thermodynamic Compensation of Urea and Trimethylamine N-Oxide Interactions with Protein. Biochemistry 1994, 33 (42), 12695–12701.

(3) Auton, M.; Bolen, D. W. Predicting the Energetics of Osmolyte-Induced Protein Folding/unfolding. Proc Natl Acad Sci U S A 2005, 102 (42), 15065–15068.

(4) Street, T. O.; Bolen, D. W.; Rose, G. D. A Molecular Mechanism for Osmolyte-Induced Protein Stability. Proc Natl Acad Sci U S A 2006, 103 (38), 13997–14002.

(5) Auton, M.; Rösgen, J.; Sinev, M.; Holthauzen, L. M. F.; Bolen, D. W. Osmolyte Effects on Protein Stability and Solubility: A Balancing Act between Backbone and Side-Chains. Biophys Chem 2011, 159 (1), 90–99.

(6) Auton, M.; Bolen, D. W.; Rösgen, J. Structural Thermodynamics of Protein Preferential Solvation: Osmolyte Solvation of Proteins, Aminoacids, and Peptides. Proteins 2008, 73 (4), 802–813.

(7) Khan, S.; Siraj, S.; Shahid, M.; Haque, M. M.; Islam, A. Osmolytes: Wonder Molecules to Combat Protein Misfolding against Stress Conditions. Int J Biol Macromol 2023, 234, 123662.

(8) Bolen, D. W.; Baskakov, I. V. The Osmophobic Effect: Natural Selection of a Thermodynamic Force in Protein Folding. J. Mol. Biol. 2001, 310 (5), 955–963.

(9) Peng, X.; Baxa, M.; Faruk, N.; Sachleben, J. R.; Pintscher, S.; Gagnon, I. A.; Houliston, S.; Arrowsmith, C. H.; Freed, K. F.; Rocklin, G. J.; Sosnick, T. R. Prediction and Validation of a Protein’s Free Energy Surface Using Hydrogen Exchange and (Importantly) Its Denaturant Dependence. J Chem Theory Comput 2022, 18 (1), 550–561.

(10) Timasheff, S. N. Solvent Stabilization of Protein Structure. Methods Mol. Biol. 1995, 40, 269–.

(11) Rösgen, J.; Pettitt, B. M.; Bolen, D. W. Protein Folding, Stability, and Solvation Structure in Osmolyte Solutions. Biophys J 2005, 89 (5), 2988–2997.

(12) Piccoli, V.; Martínez, L. Ionic Liquid Solvation of Proteins in Native and Denatured States. J. Mol. Liq. 2022, 363 (119953), 119953.

(13) Candotti, M.; Esteban-Martín, S.; Salvatella, X.; Orozco, M. Toward an Atomistic Description of the Urea-Denatured State of Proteins. Proc Natl Acad Sci U S A 2013, 110 (15), 5933–5938.

(14) Hua, L.; Zhou, R.; Thirumalai, D.; Berne, B. J. Urea Denaturation by Stronger Dispersion Interactions with Proteins than Water Implies a 2-Stage Unfolding. Proc Natl Acad Sci U S A 2008, 105 (44), 16928–16933.

(15) Rydeen, A. E.; Brustad, E. M.; Pielak, G. J. Osmolytes and Protein-Protein Interactions. J. Am. Chem. Soc. 2018, 140 (24), 7441–7444.

(16) Weerasinghe, S.; Smith, P. E. A Kirkwood−Buff Derived Force Field for Mixtures of Urea and Water. J. Phys. Chem. B 2003, 107 (16), 3891–3898.

(17) Ganguly, P.; Boserman, P.; van der Vegt, N. F. A.; Shea, J.-E. Trimethylamine N-Oxide Counteracts Urea Denaturation by Inhibiting Protein–Urea Preferential Interaction. 2017. 10.1021/jacs.7b11695.

(18) Kokubo, H.; Hu, C. Y.; Pettitt, B. M. Peptide Conformational Preferences in Osmolyte Solutions: Transfer Free Energies of Decaalanine. J Am Chem Soc 2011, 133 (6), 1849–1858.

(19) Folberth, A.; van der Vegt, N. F. A. Influence of TMAO and Pressure on the Folding Equilibrium of TrpCage. J Phys Chem B 2022, 126 (42), 8374–8380.

(20) Olgenblum, G. I.; Carmon, N.; Harries, D. Not Always Sticky: Specificity of Protein Stabilization by Sugars Is Conferred by Protein-Water Hydrogen Bonds. J Am Chem Soc 2023, 145 (42), 23308–23320.

(21) Stumpe, M. C.; Grubmüller, H. Interaction of Urea with Amino Acids: Implications for Urea-Induced Protein Denaturation. J Am Chem Soc 2007, 129 (51), 16126–16131.

(22) Lin, B.; Pettitt, B. M. Note: On the Universality of Proximal Radial Distribution Functions of Proteins. J Chem Phys 2011, 134 (10), 106101.

(23) Nguyen, B. L.; Pettitt, B. M. Effects of Acids, Bases, and Heteroatoms on Proximal Radial Distribution Functions for Proteins. J Chem Theory Comput 2015, 11 (4), 1399–1409.

(24) Martínez, L.; Shimizu, S. Molecular Interpretation of Preferential Interactions in Protein Solvation: A Solvent-Shell Perspective by Means of Minimum-Distance Distribution Functions. J. Chem. Theory Comput. 2017, 13 (12), 6358–6372.

(25) Martínez, L. ComplexMixtures.jl: Investigating the Structure of Solutions of Complex-Shaped Molecules from a Solvent-Shell Perspective. J. Mol. Liq. 2022, 347 (117945), 117945.

(26) Pereira, A. F.; Piccoli, V.; Martínez, L. Trifluoroethanol Direct Interactions with Protein Backbones Destabilize α-Helices. J. Mol. Liq. 2022, 365 (120209), 120209.

(27) Noel, J. K.; Onuchic, J. N. The Many Faces of Structure-Based Potentials: From Protein Folding Landscapes to Structural Characterization of Complex Biomolecules. In Computational Modeling of Biological Systems; Biological and Medical Physics, Biomedical Engineering; Springer US: Boston, MA, 2012; pp 31–54.

(28) Xu, W.; Harrison, S. C.; Eck, M. J. Three-Dimensional Structure of the Tyrosine Kinase c-Src. Nature 1997, 385 (6617), 595–602.

(29) Gouda, H.; Torigoe, H.; Saito, A.; Sato, M.; Arata, Y.; Shimada, I. Three-Dimensional Solution Structure of the B Domain of Staphylococcal Protein A: Comparisons of the Solution and Crystal Structures. Biochemistry 1992, 31 (40), 9665–9672.

(30) Onuchic, J. N.; Nymeyer, H.; García, A. E.; Chahine, J.; Socci, N. D. The Energy Landscape Theory of Protein Folding: Insights into Folding Mechanisms and Scenarios. Adv. Protein Chem. 2000, 53, 152–.

(31) Onuchic, J. N.; Wolynes, P. G. Theory of Protein Folding. Curr. Opin. Struct. Biol. 2004, 14 (1), 70–75.

(32) Lammert, H.; Schug, A.; Onuchic, J. N. Robustness and Generalization of Structure-Based Models for Protein Folding and Function. Proteins 2009, 77 (4), 881–891.

(33) Noel, J. K.; Schug, A.; Verma, A.; Wenzel, W.; Garcia, A. E.; Onuchic, J. N. Mirror Images as Naturally Competing Conformations in Protein Folding. J. Phys. Chem. B 2012, 116 (23), 6880–6888.

(34) Whitford, P. C.; Noel, J. K.; Gosavi, S.; Schug, A.; Sanbonmatsu, K. Y.; Onuchic, J. N. An All-Atom Structure-Based Potential for Proteins: Bridging Minimal Models with All-Atom Empirical Forcefields. Proteins 2009, 75 (2), 430–441.

(35) Oliveira, A. B., Jr; Yang, H.; Whitford, P. C.; Leite, V. B. P. Distinguishing Biomolecular Pathways and Metastable States. J. Chem. Theory Comput. 2019, 15 (11), 6482–6490.

(36) Noel, J. K.; Whitford, P. C.; Onuchic, J. N. The Shadow Map: A General Contact Definition for Capturing the Dynamics of Biomolecular Folding and Function. J. Phys. Chem. B 2012, 116 (29), 8692–8702.

(37) Demakis, C.; Childers, M. C.; Daggett, V. Conserved Patterns and Interactions in the Unfolding Transition State across SH3 Domain Structural Homologues. Protein Sci 2021, 30 (2), 391–407.

(38) Borreguero, J. M.; Ding, F.; Buldyrev, S. V.; Stanley, H. E.; Dokholyan, N. V. Multiple Folding Pathways of the SH3 Domain. Biophys J 2004, 87 (1), 521–533.

(39) Borreguero, J. M.; Dokholyan, N. V.; Buldyrev, S. V.; Shakhnovich, E. I.; Stanley, H. E. Thermodynamics and Folding Kinetics Analysis of the SH3 Domain Form Discrete Molecular Dynamics. J Mol Biol 2002, 318 (3), 863–876.

(40) Gsponer, J.; Caflisch, A. Molecular Dynamics Simulations of Protein Folding from the Transition State. Proc Natl Acad Sci U S A 2002, 99 (10), 6719–6724.

(41) Tsai, J.; Levitt, M.; Baker, D. Hierarchy of Structure Loss in MD Simulations of Src SH3 Domain Unfolding. J Mol Biol 1999, 291 (1), 215–225.

(42) Pereira, A. F.; Martínez, L. Helical Content Correlations and Hydration Structures of the Folding Ensemble of the B Domain of Protein A. J. Chem. Inf. Model. 2024, 64 (8), 3350–3359.

(43) Contessoto, V. G.; Ferreira, P. H. B.; Chahine, J.; Leite, V. B. P.; Oliveira, R. J. Small Neutral Crowding Solute Effects on Protein Folding Thermodynamic Stability and Kinetics. J Phys Chem B 2021, 125 (42), 11673–11686.

(44) Viegas, R. G.; Martins, I. B. S.; Sanches, M. N.; Oliveira Junior, A. B.; Camargo, J. B. de; Paulovich, F. V.; Leite, V. B. P. ELViM: Exploring Biomolecular Energy Landscapes through Multidimensional Visualization. J Chem Inf Model 2024, 64 (8), 3443–3450.

(45) García, A. E.; Onuchic, J. N. Folding a Protein in a Computer: An Atomic Description of the Folding/unfolding of Protein A. Proc Natl Acad Sci U S A 2003, 100 (24), 13898–13903.

(46) Khalili, M.; Liwo, A.; Scheraga, H. A. Kinetic Studies of Folding of the B-Domain of Staphylococcal Protein A with Molecular Dynamics and a United-Residue (UNRES) Model of Polypeptide Chains. J Mol Biol 2006, 355 (3), 536–547.

(47) Zou, Q.; Bennion, B. J.; Daggett, V.; Murphy, K. P. The Molecular Mechanism of Stabilization of Proteins by TMAO and Its Ability to Counteract the Effects of Urea. J Am Chem Soc 2002, 124 (7), 1192–1202.

(48) Nasralla, M.; Laurent, H.; Alderman, O. L. G.; Headen, T. F.; Dougan, L. Trimethylamine-N-Oxide Depletes Urea in a Peptide Solvation Shell. Proceedings of the National Academy of Sciences 2024, 121 (14), e2317825121.

(49) Timasheff, S. N. Protein-Solvent Preferential Interactions, Protein Hydration, and the Modulation of Biochemical Reactions by Solvent Components. Proc Natl Acad Sci U S A 2002, 99 (15), 9721–9726.

(50) Auton, M.; Bolen, D. W. Predicting the Energetics of Osmolyte-Induced Protein Folding/unfolding. Proc Natl Acad Sci U S A 2005, 102 (42), 15065–15068.

(51) Street, T. O.; Bolen, D. W.; Rose, G. D. A Molecular Mechanism for Osmolyte-Induced Protein Stability. Proc Natl Acad Sci U S A 2006, 103 (38), 13997–14002.

(52) Auton, M.; Rösgen, J.; Sinev, M.; Holthauzen, L. M. F.; Bolen, D. W. Osmolyte Effects on Protein Stability and Solubility: A Balancing Act between Backbone and Side-Chains. Biophys Chem 2011, 159 (1), 90–99.

(53) Auton, M.; Holthauzen, L. M. F.; Bolen, D. W. Anatomy of Energetic Changes Accompanying Urea-Induced Protein Denaturation. Proc Natl Acad Sci U S A 2007, 104 (39), 15317–15322.

(54) Auton, M.; Bolen, D. W.; Rösgen, J. Structural Thermodynamics of Protein Preferential Solvation: Osmolyte Solvation of Proteins, Aminoacids, and Peptides. Proteins 2008, 73 (4), 802–813.

(55) Holthauzen, L. M. F.; Rösgen, J.; Bolen, D. W. Hydrogen Bonding Progressively Strengthens upon Transfer of the Protein Urea-Denatured State to Water and Protecting Osmolytes. Biochemistry 2010, 49 (6), 1310–1318.

(56) Okuno, Y.; Yoo, J.; Schwieters, C. D.; Best, R. B.; Chung, H. S.; Clore, G. M. Atomic View of Cosolute-Induced Protein Denaturation Probed by NMR Solvent Paramagnetic Relaxation Enhancement. Proc Natl Acad Sci U S A 2021, 118 (34). 10.1073/pnas.2112021118.

(57) Courtenay, E. S.; Capp, M. W.; Anderson, C. F.; Record, M. T., Jr. Vapor Pressure Osmometry Studies of Osmolyte-Protein Interactions: Implications for the Action of Osmoprotectants in Vivo and for the Interpretation of “Osmotic Stress” Experiments in Vitro. Biochemistry 2000, 39 (15), 4455–4471.

(58) Roy, S.; Bagchi, B. Comparative Study of Protein Unfolding in Aqueous Urea and Dimethyl Sulfoxide Solutions: Surface Polarity, Solvent Specificity, and Sequence of Secondary Structure Melting. J. Phys. Chem. B 2014, 118 (21), 5691–5697.

(59) Parui, S.; Jana, B. Relative Solvent Exposure of the Alpha-Helix and Beta-Sheet in Water Determines the Initial Stages of Urea and Guanidinium Chloride-Induced Denaturation of Alpha/Beta Proteins. J. Phys. Chem. B 2019, 123 (42), 8889–8900.

(60) Bennion, B. J.; Daggett, V. The Molecular Basis for the Chemical Denaturation of Proteins by Urea. Proc. Natl. Acad. Sci. U. S. A. 2003, 100 (9), 5142–5147.

(61) Wyman, J. Linked Functions and Reciprocal Effects in Hemoglobin: A Second Look. Adv. Prot. Chem. 1964, 19, 286–.

(62) Canchi, D. R.; García, A. E. Cosolvent Effects on Protein Stability. Annu. Rev. Phys. Chem. 2013, 64, 293–.

(63) Bezsonova, I.; Singer, A.; Choy, W.-Y.; Tollinger, M.; Forman-Kay, J. D. Structural Comparison of the Unstable drkN SH3 Domain and a Stable Mutant. Biochemistry 2005, 44 (47), 15550–15560.

(64) Tanford, C. Protein Denaturation. In Advances in Protein Chemistry; Advances in protein chemistry; Elsevier, 1970; pp 1–95.

(65) Tanford, C. Isothermal Unfolding of Globular Proteins in Aqueous Urea Solutions. J. Am. Chem. Soc. 1964, 86 (10), 2050–2059.

(66) Tanford, C. Contribution of Hydrophobic Interactions to the Stability of the Globular Conformation of Proteins. J. Am. Chem. Soc. 1962, 84 (22), 4240–4247.

(67) Wang, A.; Bolen, D. W. A Naturally Occurring Protective System in Urea-Rich Cells: Mechanism of Osmolyte Protection of Proteins against Urea Denaturation. Biochemistry 1997, 36 (30), 9101–9108.

(68) Auton, M.; Bolen, D. W. Additive Transfer Free Energies of the Peptide Backbone Unit That Are Independent of the Model Compound and the Choice of Concentration Scale. Biochemistry 2004, 43 (5), 1329–1342.

(69) Creamer, T. P.; Srinivasan, R.; Rose, G. D. Modeling Unfolded States of Proteins and Peptides. II. Backbone Solvent Accessibility. Biochemistry 1997, 36 (10), 2832–2835.

(70) Schellman, J. A. Protein Stability in Mixed Solvents: A Balance of Contact Interaction and Excluded Volume. Biophys J 2003, 85 (1), 108–125.

(71) Auton, M.; Bolen, D. W. Application of the Transfer Model to Understand How Naturally Occurring Osmolytes Affect Protein Stability. Methods Enzymol 2007, 428, 418–.

(72) Auton, M.; Holthauzen, L. M. F.; Bolen, D. W. Anatomy of Energetic Changes Accompanying Urea-Induced Protein Denaturation. Proc Natl Acad Sci U S A 2007, 104 (39), 15317–15322.

(73) Holthauzen, L. M. F.; Rösgen, J.; Bolen, D. W. Hydrogen Bonding Progressively Strengthens upon Transfer of the Protein Urea-Denatured State to Water and Protecting Osmolytes. Biochemistry 2010, 49 (6), 1310–1318.

(74) Ziv, G.; Haran, G. Protein Folding, Protein Collapse, and Tanford’s Transfer Model: Lessons from Single-Molecule FRET. J Am Chem Soc 2009, 131 (8), 2942–2947.

(75) Taketomi, H.; Ueda, Y.; Gō, N. Studies on Protein Folding, Unfolding and Fluctuations by Computer Simulation. I. The Effect of Specific Amino Acid Sequence Represented by Specific Inter-Unit Interactions. Int. J. Pept. Protein Res. 1975, 7 (6), 445–459.

(76) Clementi, C.; Nymeyer, H.; Onuchic, J. N. Topological and Energetic Factors: What Determines the Structural Details of the Transition State Ensemble and “En-Route” Intermediates for Protein Folding? An Investigation for Small Globular Proteins. J. Mol. Biol. 2000, 298 (5), 937–953.

(77) Sobolev, V.; Sorokine, A.; Prilusky, J.; Abola, E. E.; Edelman, M. Automated Analysis of Interatomic Contacts in Proteins. Bioinformatics 1999, 15 (4), 327–332.

(78) Noel, J. K.; Whitford, P. C.; Sanbonmatsu, K. Y.; Onuchic, J. N. SMOG@ctbp: Simplified Deployment of Structure-Based Models in GROMACS. Nucleic Acids Res. 2010, 38 (Web Server issue), W657–W661.

(79) Hess, B.; Kutzner, C.; van der Spoel, D.; Lindahl, E. GROMACS 4: Algorithms for Highly Efficient, Load-Balanced, and Scalable Molecular Simulation. J. Chem. Theory Comput. 2008, 4 (3), 435–447.

(80) Kumar, S.; Rosenberg, J. M.; Bouzida, D.; Swendsen, R. H.; Kollman, P. A. THE Weighted Histogram Analysis Method for Free-Energy Calculations on Biomolecules. I. The Method. J. Comput. Chem. 1992, 13 (8), 1011–1021.

(81) Noel, J. K.; Levi, M.; Raghunathan, M.; Lammert, H.; Hayes, R. L.; Onuchic, J. N.; Whitford, P. C. SMOG 2: A Versatile Software Package for Generating Structure-Based Models. PLoS Comput. Biol. 2016, 12 (3), e1004794.

(82) Oliveira Junior, A. B.; Lin, X.; Kulkarni, P.; Onuchic, J. N.; Roy, S.; Leite, V. B. P. Exploring Energy Landscapes of Intrinsically Disordered Proteins: Insights into Functional Mechanisms. J. Chem. Theory Comput. 2021, 17 (5), 3178–3187.

(83) Dias, R. V. R.; Pedro, R. P.; Sanches, M. N.; Moreira, G. C.; Leite, V. B. P.; Caruso, I. P.; de Melo, F. A.; de Oliveira, L. C. Unveiling Metastable Ensembles of GRB2 and the Relevance of Interdomain Communication during Folding. J. Chem. Inf. Model. 2023, 63 (20), 6344–6353.

(84) Sanches, M. N.; Parra, R. G.; Viegas, R. G.; Oliveira, A. B., Jr; Wolynes, P. G.; Ferreiro, D. U.; Leite, V. B. P. Resolving the Fine Structure in the Energy Landscapes of Repeat Proteins. QRB Discov 2022, 3, e7.

(85) Viegas, R. G.; Sanches, M. N.; Chen, A. A.; Paulovich, F. V.; Garcia, A. E.; Leite, V. B. P. Characterizing the Folding Transition-State Ensembles in the Energy Landscape of an RNA Tetraloop. J. Chem. Inf. Model. 2023, 63 (17), 5641–5649.

(86) da Silva, F. B.; Simien, J. M.; Viegas, R. G.; Haglund, E.; Leite, V. B. P. Exploring the Folding Landscape of Leptin: Insights into Threading Pathways. J. Struct. Biol. 2023, 216 (1), 108054.

(87) Hardin, C.; Eastwood, M. P.; Prentiss, M. C.; Luthey-Schulten, Z.; Wolynes, P. G. Associative Memory Hamiltonians for Structure Prediction without Homology: Alpha/beta Proteins. Proc. Natl. Acad. Sci. U. S. A. 2003, 100 (4), 1679–1684.

(88) Kabsch, W.; Sander, C. Dictionary of Protein Secondary Structure: Pattern Recognition of Hydrogen-Bonded and Geometrical Features. Biopolymers 1983, 22 (12), 2577–2637.

(89) Joosten, R. P.; te Beek, T. A. H.; Krieger, E.; Hekkelman, M. L.; Hooft, R. W. W.; Schneider, R.; Sander, C.; Vriend, G. A Series of PDB Related Databases for Everyday Needs. Nucleic Acids Res. 2011, 39 (Database issue), D411–D419.

(90) Rotkiewicz, P.; Skolnick, J. Fast Procedure for Reconstruction of Full-Atom Protein Models from Reduced Representations. J. Comput. Chem. 2008, 29 (9), 1460–1465.

(91) Martínez, L.; Andrade, R.; Birgin, E. G.; Martínez, J. M. PACKMOL: A Package for Building Initial Configurations for Molecular Dynamics Simulations. J. Comput. Chem. 2009, 30 (13), 2157–2164.

(92) Martínez, J. M.; Martínez, L. Packing Optimization for Automated Generation of Complex System’s Initial Configurations for Molecular Dynamics and Docking. J. Comput. Chem. 2003, 24 (7), 819–825.

(93) MacKerell, A. D.; Bashford, D.; Bellott, M.; Dunbrack, R. L.; Evanseck, J. D.; Field, M. J.; Fischer, S.; Gao, J.; Guo, H.; Ha, S.; Joseph-McCarthy, D.; Kuchnir, L.; Kuczera, K.; Lau, F. T.; Mattos, C.; Michnick, S.; Ngo, T.; Nguyen, D. T.; Prodhom, B.; Reiher, W. E.; Roux, B.; Schlenkrich, M.; Smith, J. C.; Stote, R.; Straub, J.; Watanabe, M.; Wiórkiewicz-Kuczera, J.; Yin, D.; Karplus, M. All-Atom Empirical Potential for Molecular Modeling and Dynamics Studies of Proteins. J. Phys. Chem. B 1998, 102 (18), 3586–3616.

(94) Jorgensen, W. L.; Chandrasekhar, J.; Madura, J. D.; Impey, R. W.; Klein, M. L. Comparison of Simple Potential Functions for Simulating Liquid Water. The Journal of Chemical Physics. 1983, pp 926–935. 10.1063/1.445869.

(95) Vanommeslaeghe, K.; Hatcher, E.; Acharya, C.; Kundu, S.; Zhong, S.; Shim, J.; Darian, E.; Guvench, O.; Lopes, P.; Vorobyov, I.; Mackerell, A. D. CHARMM General Force Field: A Force Field for Drug-like Molecules Compatible with the CHARMM All-Atom Additive Biological Force Fields. Journal of Computational Chemistry. 2009, p NA – NA. 10.1002/jcc.21367.

(96) Yu, W.; He, X.; Vanommeslaeghe, K.; MacKerell, A. D. Extension of the CHARMM General Force Field to Sulfonyl-Containing Compounds and Its Utility in Biomolecular Simulations. Journal of Computational Chemistry. 2012, pp 2451–2468. 10.1002/jcc.23067.

(97) Schneck, E.; Horinek, D.; Netz, R. R. Insight into the Molecular Mechanisms of Protein Stabilizing Osmolytes from Global Force-Field Variations. J Phys Chem B 2013, 117 (28), 8310–8321.

(98) Ganguly, P.; Bubák, D.; Polák, J.; Fagan, P.; Dračínský, M.; van der Vegt, N. F. A.; Heyda, J.; Shea, J.-E. Cosolvent Exclusion Drives Protein Stability in Trimethylamine -Oxide and Betaine Solutions. J Phys Chem Lett 2022, 13 (34), 7980–7986.

(99) Van Der Spoel, D.; Lindahl, E.; Hess, B.; Groenhof, G.; Mark, A. E.; Berendsen, H. J. C. GROMACS: Fast, Flexible, and Free. J. Comput. Chem. 2005, 26 (16), 1701–1718.

(100) Berendsen, H. J. C.; Postma, J. P. M.; van Gunsteren, W. F.; DiNola, A.; Haak, J. R. Molecular Dynamics with Coupling to an External Bath. J. Chem. Phys. 1984, 81 (8), 3684–3690.

(101) Parrinello, M.; Rahman, A. Polymorphic Transitions in Single Crystals: A New Molecular Dynamics Method. J. Appl. Phys. 1981, 52 (12), 7182–7190.

(102) Newman, K. E. Kirkwood–Buff Solution Theory: Derivation and Applications. Chem. Soc. Rev. 1994, 23 (1), 31–40.

(103) Piccoli, V.; Martínez, L. Correlated Counterion Effects on the Solvation of Proteins by Ionic Liquids. J. Mol. Liq. 2020, 320 (114347), 114347.

(104) Christ, S.; Schwabeneder, D.; Rackauckas, C.; Borregaard, M. K.; Breloff, T. Plots.Jl – A User Extendable Plotting API for the Julia Programming Language. J. Open Res. Softw. 2023, 11. 10.5334/jors.431.

