## Supplementary Material for "Osmolyte structural and thermodynamic effects across the protein folding landscape"

**Contents:**

1. Structural characterization of folding ensembles (Table S1)
2. SBM folds space characterization (Figures S1 and S2)
3. Representative structure of each fold subset (Figure S3)
4. Secondary structure content per residue in each subset (Figures S4 and S5)
5. Correlation between Γ and SASA (Figure S6)
6. MDDF decompositions of urea and TMAO by backbone and side chain contributions (Figure S7)
7. Density maps for water (Figures S8 and S11)
8. MDDFs for all ensembles for SH3 and TMAO (Figure S9) and group contributions (Figure S10)
9. Residue contributions differential density maps for all states in urea and TMAO (Figures S12 to S15)
10. Γ for selected folds at multiple concentrations (Figure S16)
11. Simulation box details (Tables S2 to S5).

**Table S1**. Fraction of native contacts (Q), Solvent Accessible Surface Area (SASA), secondary structure content (β-sheet (SH3) and $\alpha$-helix (BdpA)), preferential interaction parameters of Urea and TMAO for each ensemble of the SH3 and BdpA proteins.

| SH3 protein | | | | | |
|---|---|---|---|---|---|
| Ensembles | Average Q | Average SASA ($nm^2$) | Average β-sheet content | Average $\Gamma_{TMAO}$ | Average $\Gamma_{Urea}$ |
| $N_{80}^{S0}$ | 0.850 | 42.413 | 80.018 | -2.416 | 2.800 |
| $U_{46}^{S1}$ | 0.553 | 48.828 | 45.584 | -2.737 | 3.185 |
| $U_{43}^{S2}$ | 0.552 | 47.512 | 42.650 | -2.752 | 3.309 |
| $U_{33}^{S3}$ | 0.544 | 47.831 | 33.044 | -2.865 | 3.659 |
| $U_{6.0}^{S4}$ | 0.236 | 56.175 | 5.999 | -3.175 | 3.846 |
| $U_{2.4}^{S5}$ | 0.176 | 58.718 | 2.437 | -3.309 | 4.355 |
| $U_{3.9}^{S6}$ | 0.170 | 62.640 | 3.932 | -3.464 | 4.519 |
| $U_{2.3}^{S7}$ | 0.159 | 62.128 | 2.258 | -3.391 | 4.260 |
| $U_{2.2}^{S8}$ | 0.139 | 65.246 | 2.220 | -3.509 | 4.258 |
| $U_{0.9}^{S9}$ | 0.106 | 67.949 | 0.892 | -3.783 | 4.661 |
| $U_{0.7}^{S10}$ | 0.097 | 66.900 | 0.689 | -3.575 | 4.617 |
| $U_{0.5}^{S11}$ | 0.091 | 68.472 | 0.502 | -3.645 | 4.651 |
| $U_{1.0}^{S12}$ | 0.087 | 69.213 | 1.040 | -3.642 | 4.860 |
| BdpA protein | | | | | |
| Ensembles | Average Q | Average SASA ($nm^2$) | Average $\alpha$-helix content | Average $\Gamma_{TMAO}$ | Average $\Gamma_{Urea}$ |
| $N_{84}^{B0}$ | 0.772 | 51.248 | 84.495 | -3.198 | 2.685 |
| $U_{63}^{B1}$ | 0.729 | 52.156 | 62.579 | -3.219 | 2.731 |
| $U_{40}^{B2}$ | 0.609 | 53.069 | 40.224 | -3.382 | 2.702 |
| $U_{38}^{B3}$ | 0.553 | 53.303 | 37.951 | -3.496 | 3.014 |
| $U_{39}^{B4}$ | 0.543 | 52.673 | 38.889 | -3.346 | 2.755 |
| $U_{37}^{B5}$ | 0.378 | 58.953 | 36.962 | -3.579 | 3.189 |
| $U_{36}^{B6}$ | 0.370 | 57.210 | 35.996 | -3.461 | 3.337 |
| $U_{15}^{B7}$ | 0.348 | 59.481 | 15.067 | -3.675 | 3.327 |
| $U_{17}^{B8}$ | 0.336 | 61.114 | 17.224 | -3.616 | 3.281 |
| $U_{14}^{B9}$ | 0.335 | 59.255 | 13.642 | -3.685 | 3.077 |
| $U_{38}^{B10}$ | 0.333 | 59.279 | 37.873 | -3.631 | 3.520 |
| $U_{37}^{B11}$ | 0.305 | 64.460 | 36.861 | -3.874 | 3.571 |

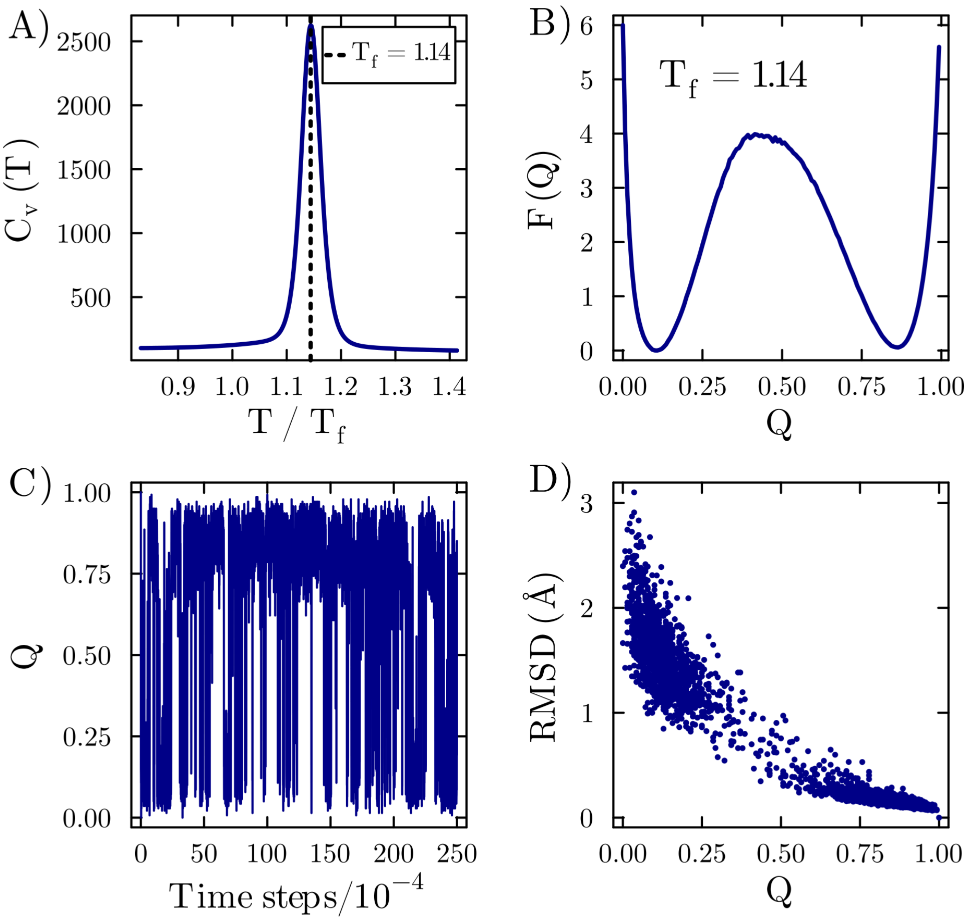


**Figure S1**. Characterization of SH3 domain folding. A) Specific heat ($C_v$) as a function of temperature, allowing the folding temperature identification ($T_f$ = 0.97 reduced units). From the simulation performed at the $T_f$: B) Free energy as a function of the fraction of native contacts (Q). C) Fraction of native contacts (Q) as a function of the simulation time step. D) Contour maps of the Probability Density (PD) as a function of Q and RMSD.

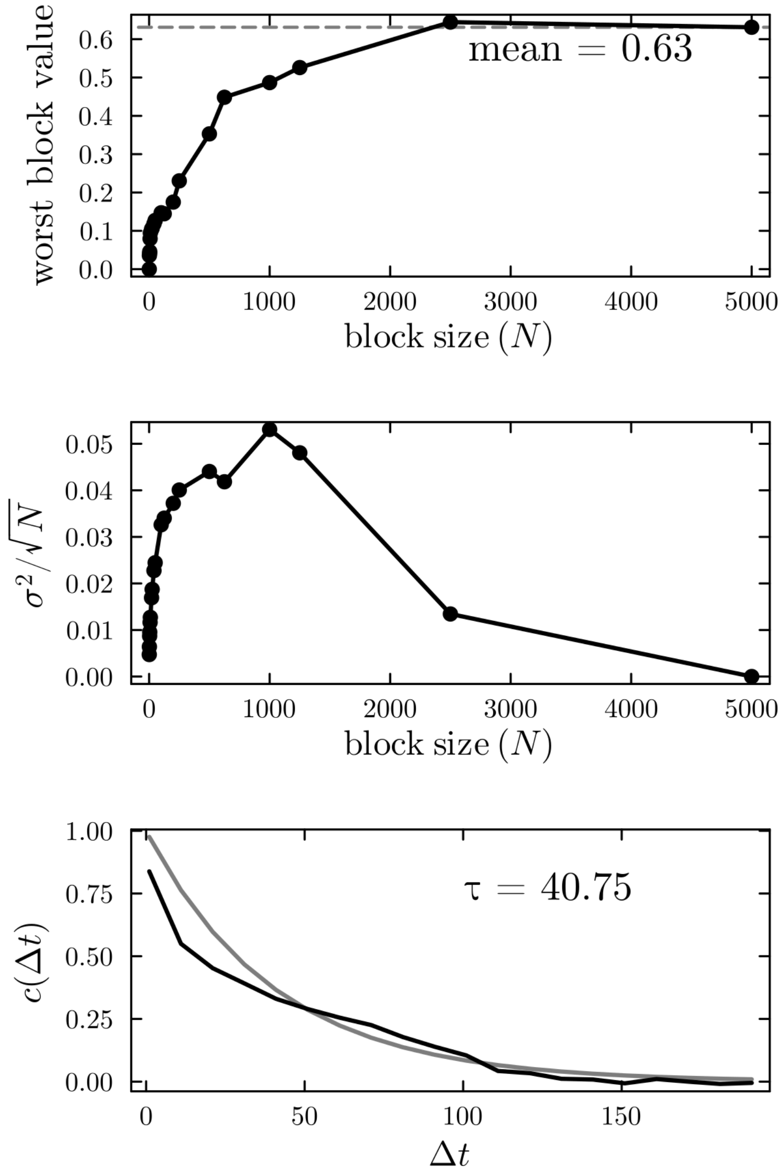


**Figure S2**. Block-averaging and autocorrelation function for the fraction of native contacts (Q) derived from simulation with $C_\alpha$-SBMs at the folding temperature ($T_f$). The analysis employed the block_average function of the MolSimToolkit.jl package, accessible at http://github.com/m3g/MolSimToolkit.jl/ - version 1.3.4. The presented data distinctly show simulation convergence, as the block average value rapidly approaches the average value of the property (Q), the standard deviation is low, and the correlation time decays quickly.

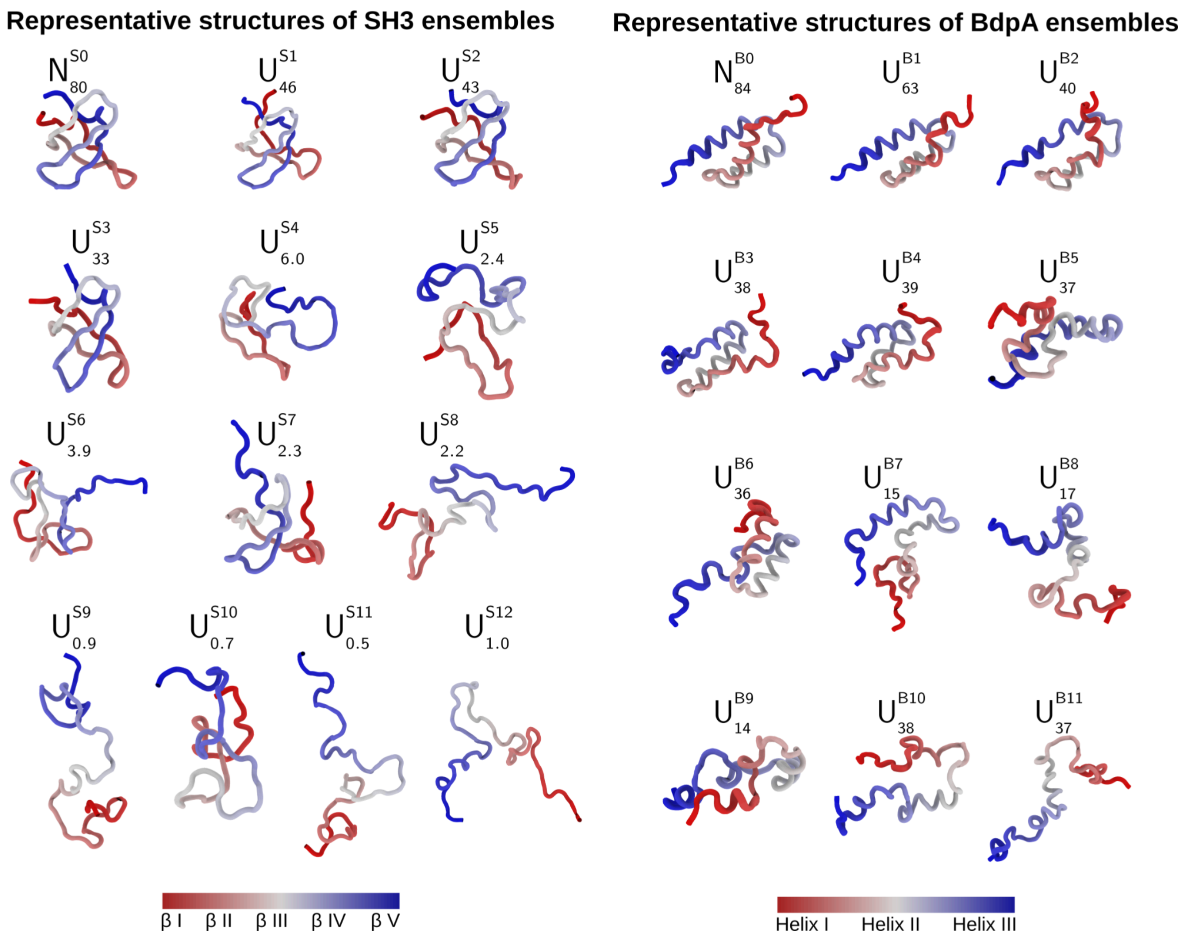


**Figure S3**. Representative structures of SH3 and BdpA ensembles.

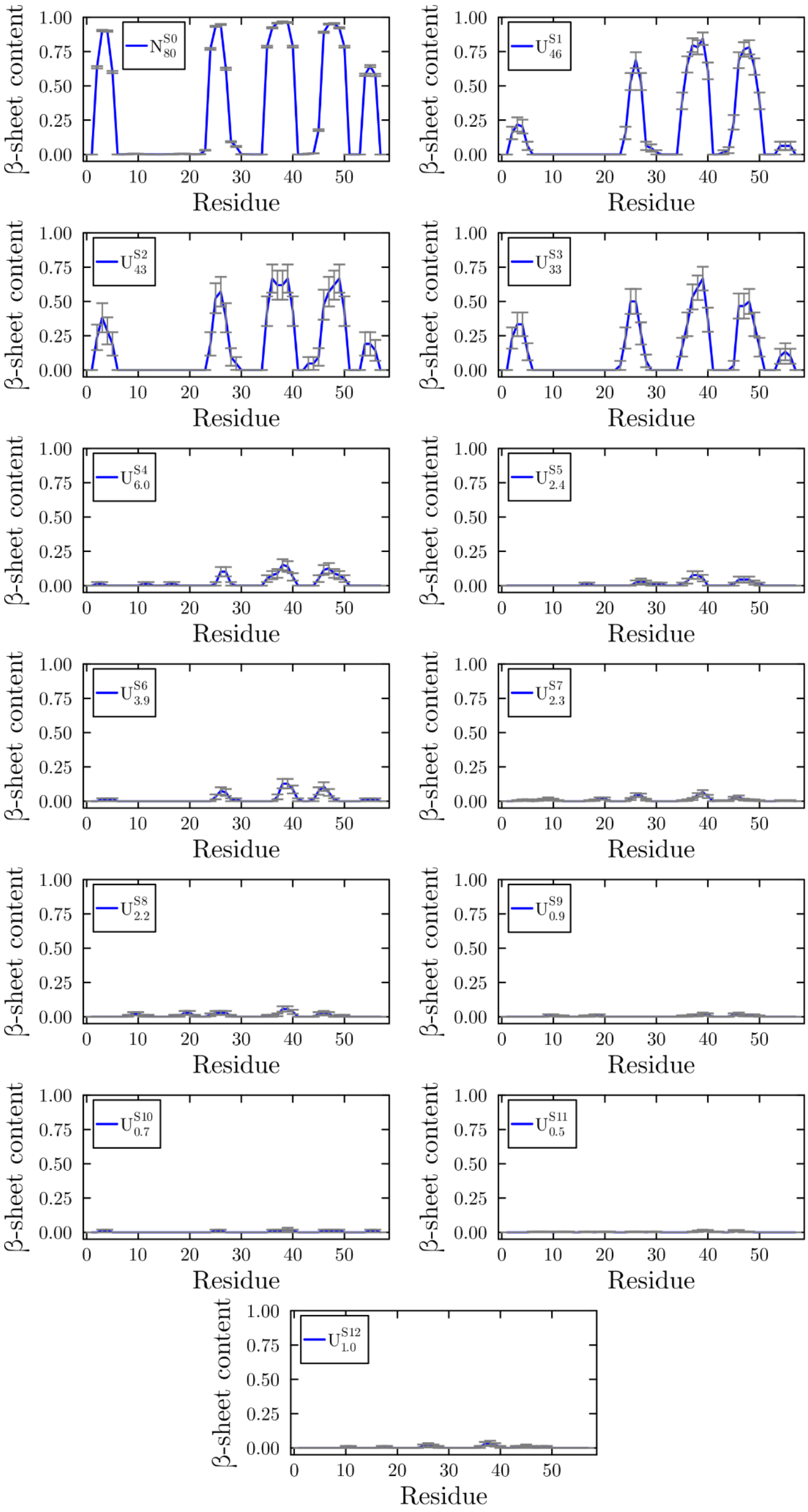


**Figure S4**. $\beta$-sheet content per residue in each SH3 folding subset.

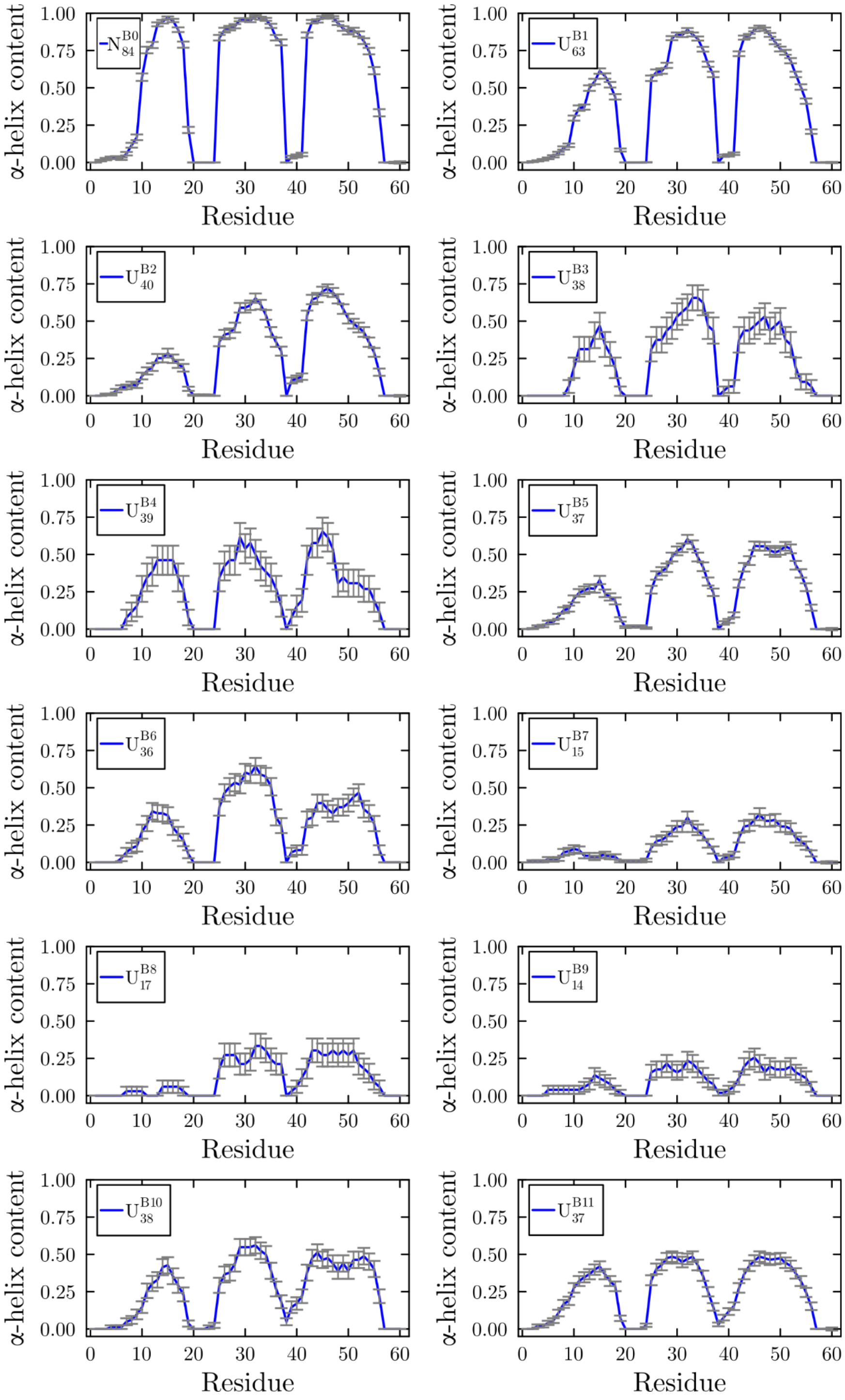


**Figure S5**. Helical content per residue in each BdpA folding subset.

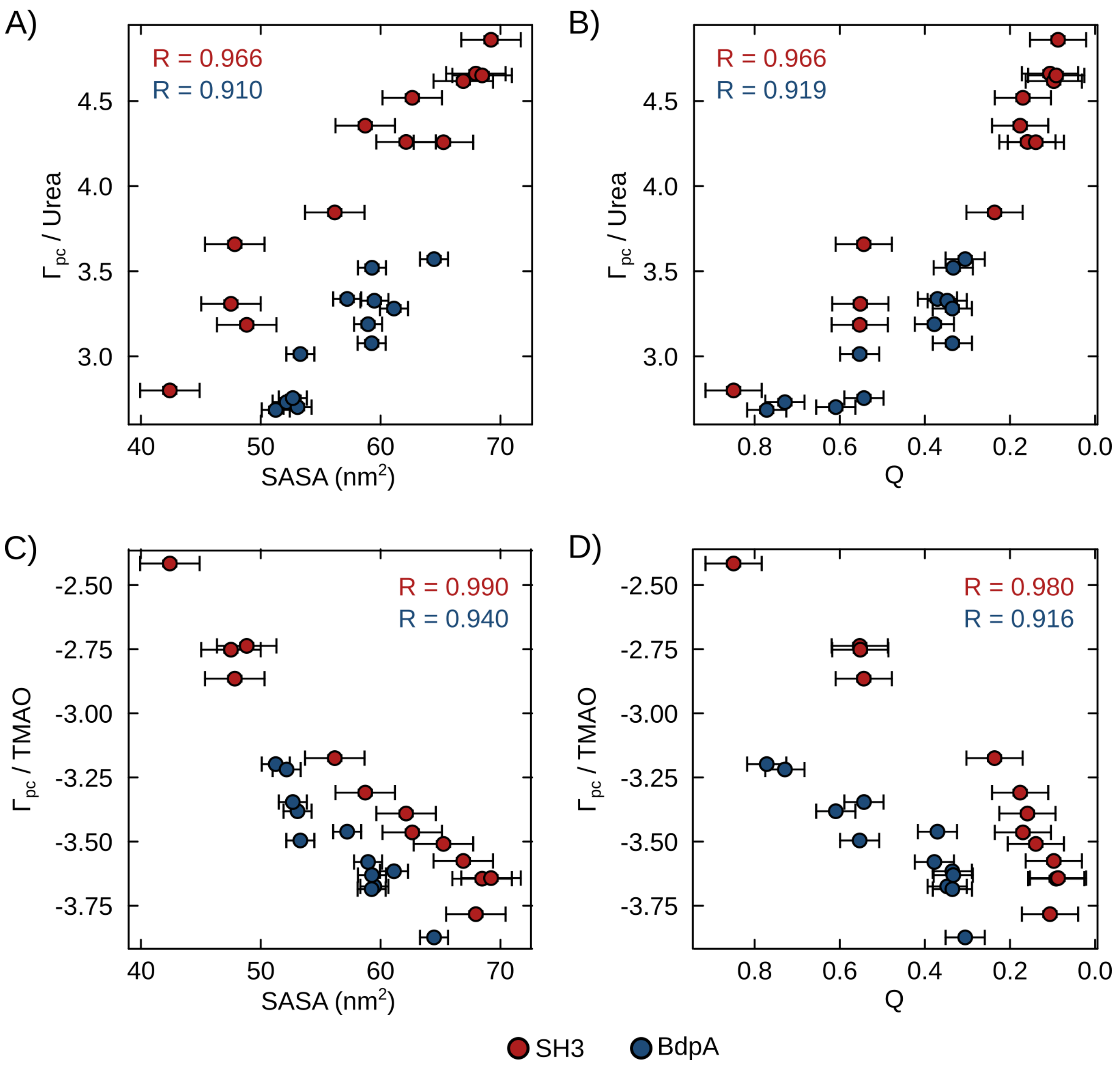


**Figure S6**. Preferential integration parameters (Γs) as a function of SASA (A) and C)) and Q (B) and D)). Red and blue dots represent the average parameters for SH3 and BdpA protein ensembles, respectively, with corresponding error bars. A quadratic fit effectively captures the correlation between Γs and SASA or Q in each case. For SH3, Q is strongly correlated with both SASA (R = 0.984) and β-sheet content (R = 0.993). For BdpA, Q also shows strong correlations with SASA (R = 0.962) and *α*-helix content (R = 0.891).

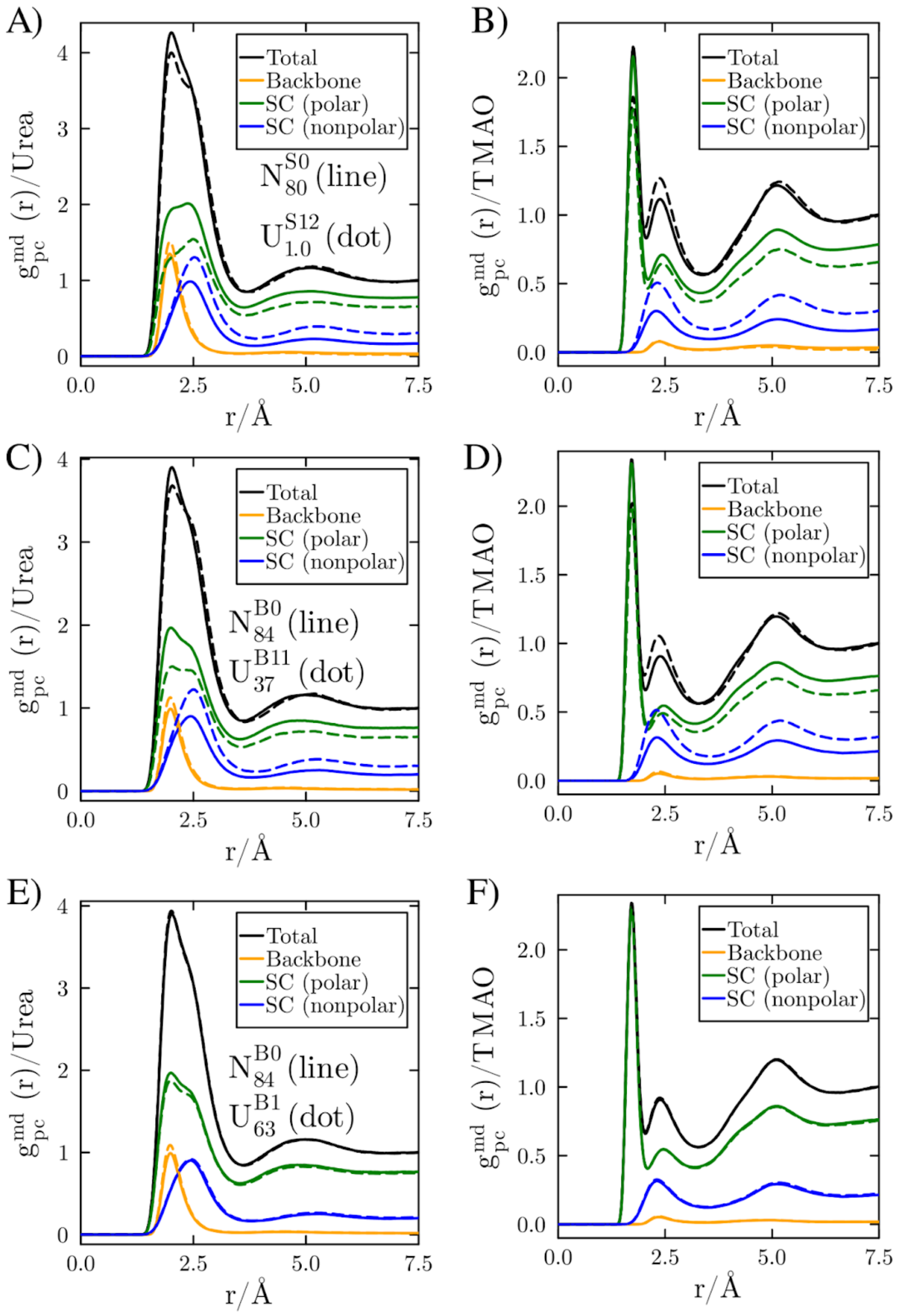


**Figure S7**. Decomposition of the MDDFs of urea (panels A, C, and E) and TMAO (panels B, D, and F) into contributions from the backbone (orange), polar side chain (green), and nonpolar side chain (blue). Solid lines represent the native ensembles, and dashed lines represent the denatured ensembles. Panels A and B compare the SH3 domain ensembles $N_{80}^{S0}$ and $U_{1.0}^{S12}$ in urea and TMAO, respectively. Panels C and D show MDDFs for BdpA, comparing ensembles $N_{84}^{B0}$ and $U_{37}^{B11}$ in urea and TMAO, respectively. Panels E and F show the same comparison as in C and D, but with a partially denatured ensemble $U_{63}^{B1}$.

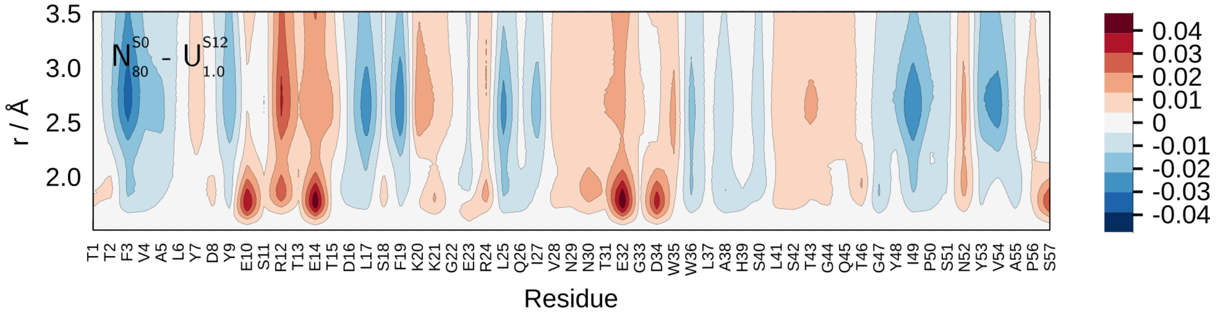


**Figure S8**. Difference in the MDDF density of the water in the vicinity of $N_{80}^{S0}$ and unfolded ($U_{1.0}^{S12}$) state in the urea 0.5 mol L$^{-1}$ solution. Red regions indicate higher water density near the $N_{80}^{S0}$ state, while blue regions show higher density around the $U_{1.0}^{S12}$ state. This pattern emphasizes the increased interactions of the solvent with mostly hydrophobic residues in the $U_{1.0}^{S12}$ state that are typically protected from the solvent in the $N_{80}^{S0}$ conformation.

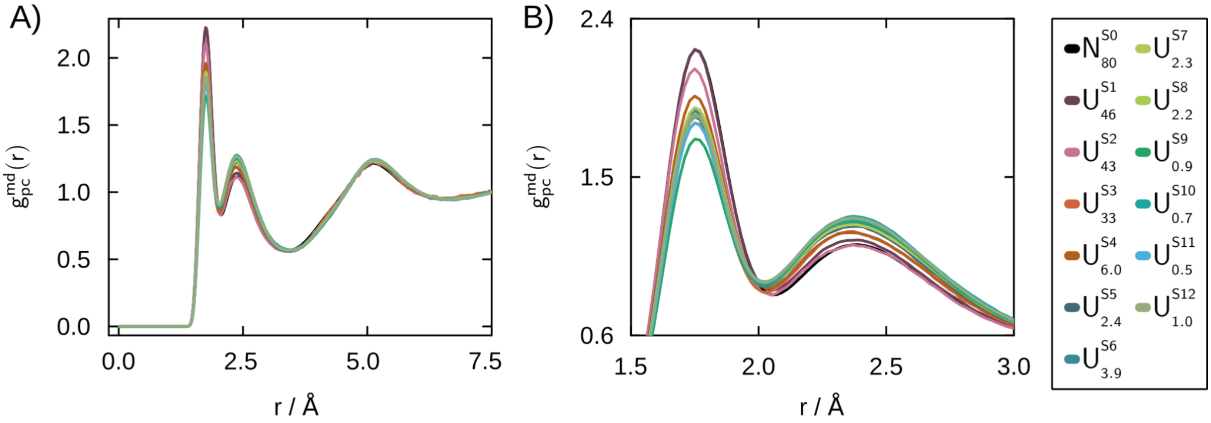


**Figure S9**. MDDF of TMAO 0.5 mol $L^{-1}$ for all ensembles of the SH3 protein.

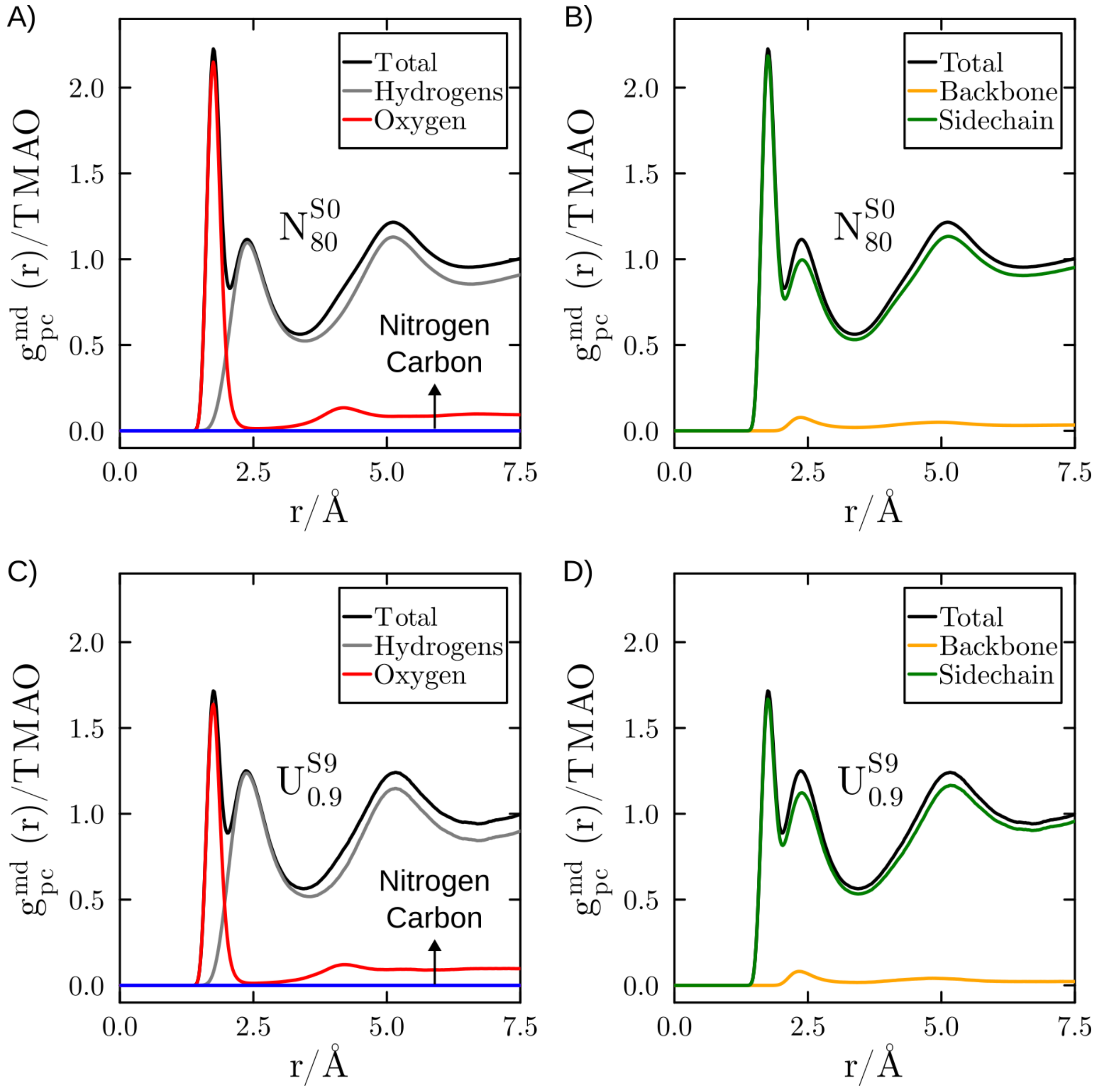


**Figure S10**. Total MDDF of TMAO and group contributions for the $N_{80}^{S0}$ and $U_{0.9}^{S9}$ ensembles. A) and C) show the contributions of the atoms and atom groups of the TMAO's MDDF of $N_{80}^{S0}$ and $U_{0.9}^{S9}$ ensembles. B) and D) show the respective contributions of the backbone (yellow) and side chain (green) to the total TMAO's MDDF.

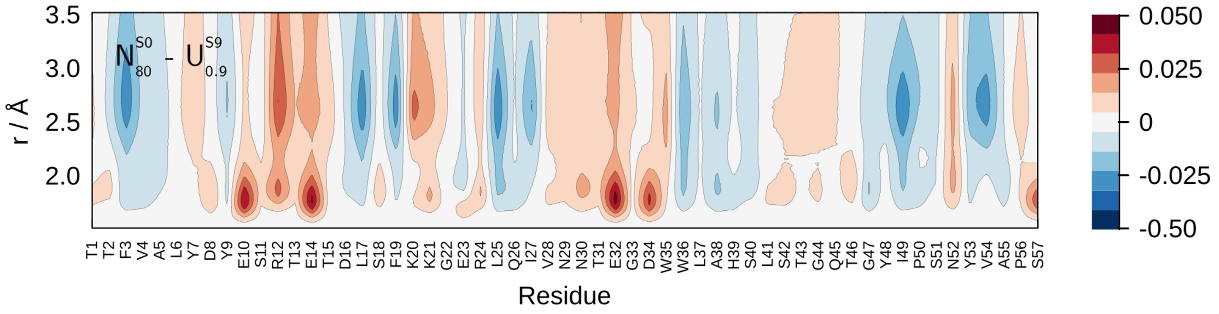


**Figure S11**. Difference in the MDDF density of the water in the vicinity of $N_{80}^{S0}$ and unfolded ($U_{0.9}^{S9}$) state in the TMAO 0.5 mol L$^{-1}$ solution. Red regions indicate higher water density near the $N_{80}^{S0}$ state, while blue regions show higher density around the $U_{0.9}^{S9}$ state. This pattern emphasizes the increased interactions of the solvent with mostly hydrophobic residues in the $U_{0.9}^{S9}$ state that are typically protected from the solvent in the $N_{80}^{S0}$ conformation.

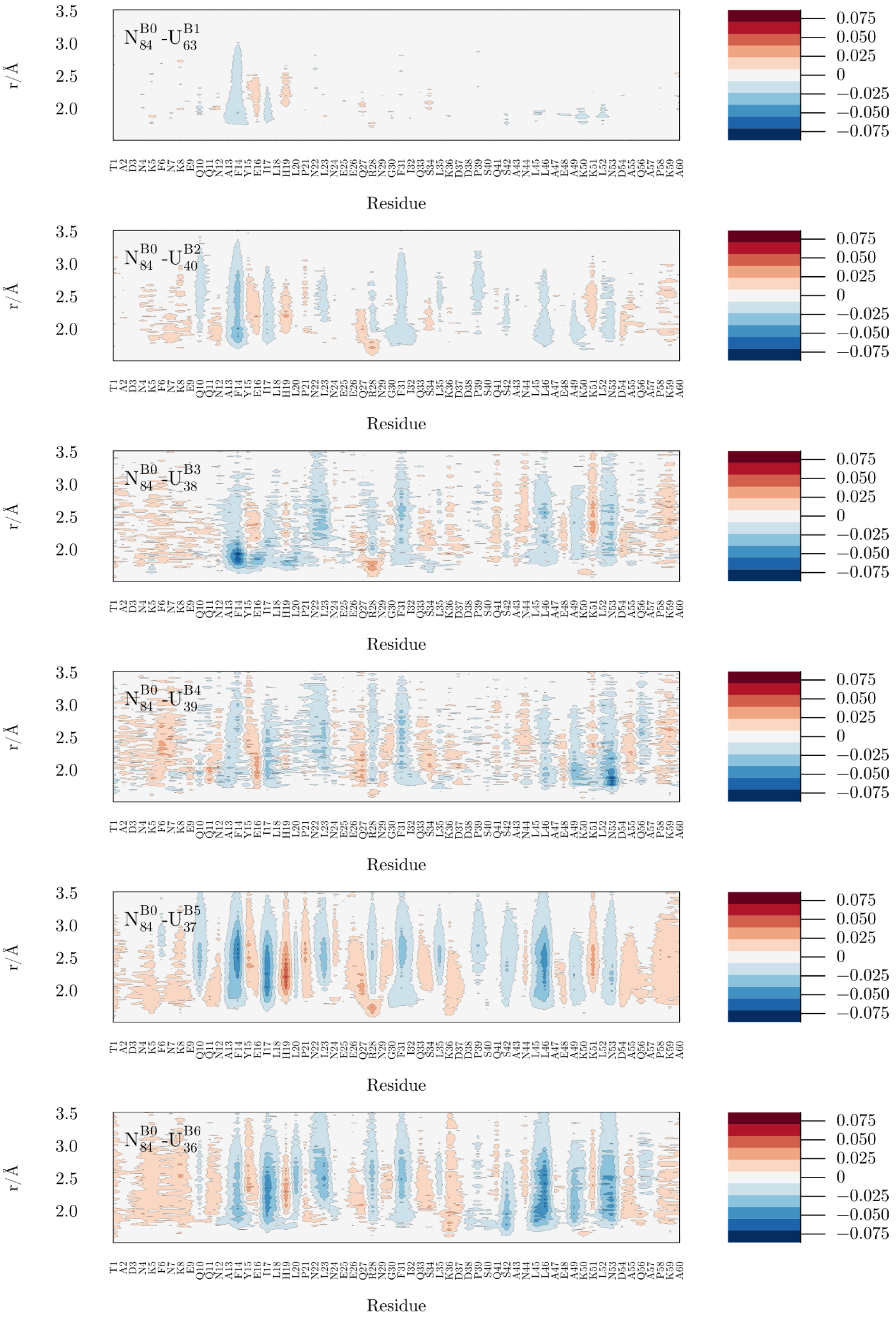


**Figure S12**. Differential density maps per residue for BdpA in urea solution 0.5 mol L$^{-1}$.

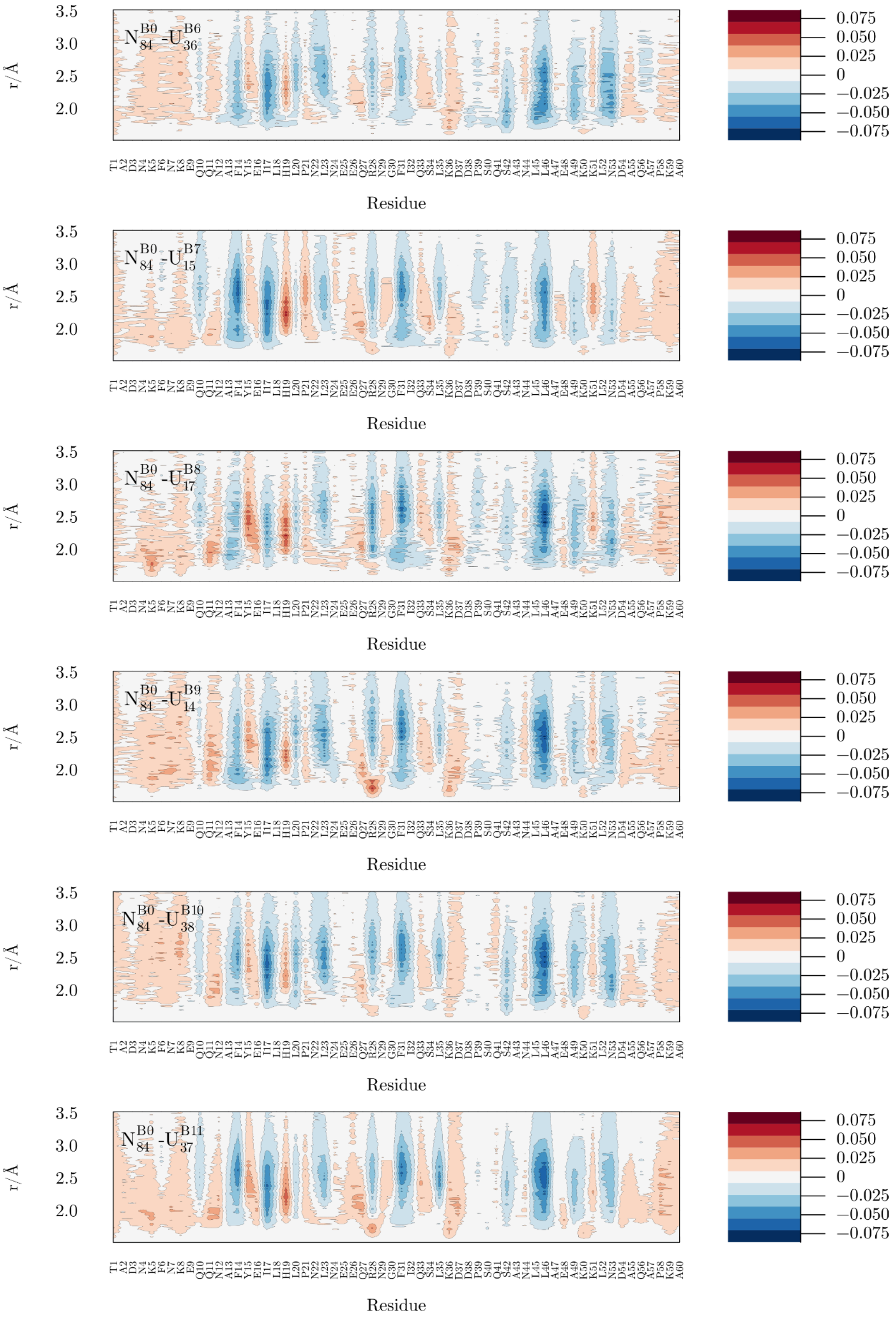


**Figure S12 (continued)**. Differential density maps per residue for BdpA in urea solution 0.5 mol $L^{-1}$.

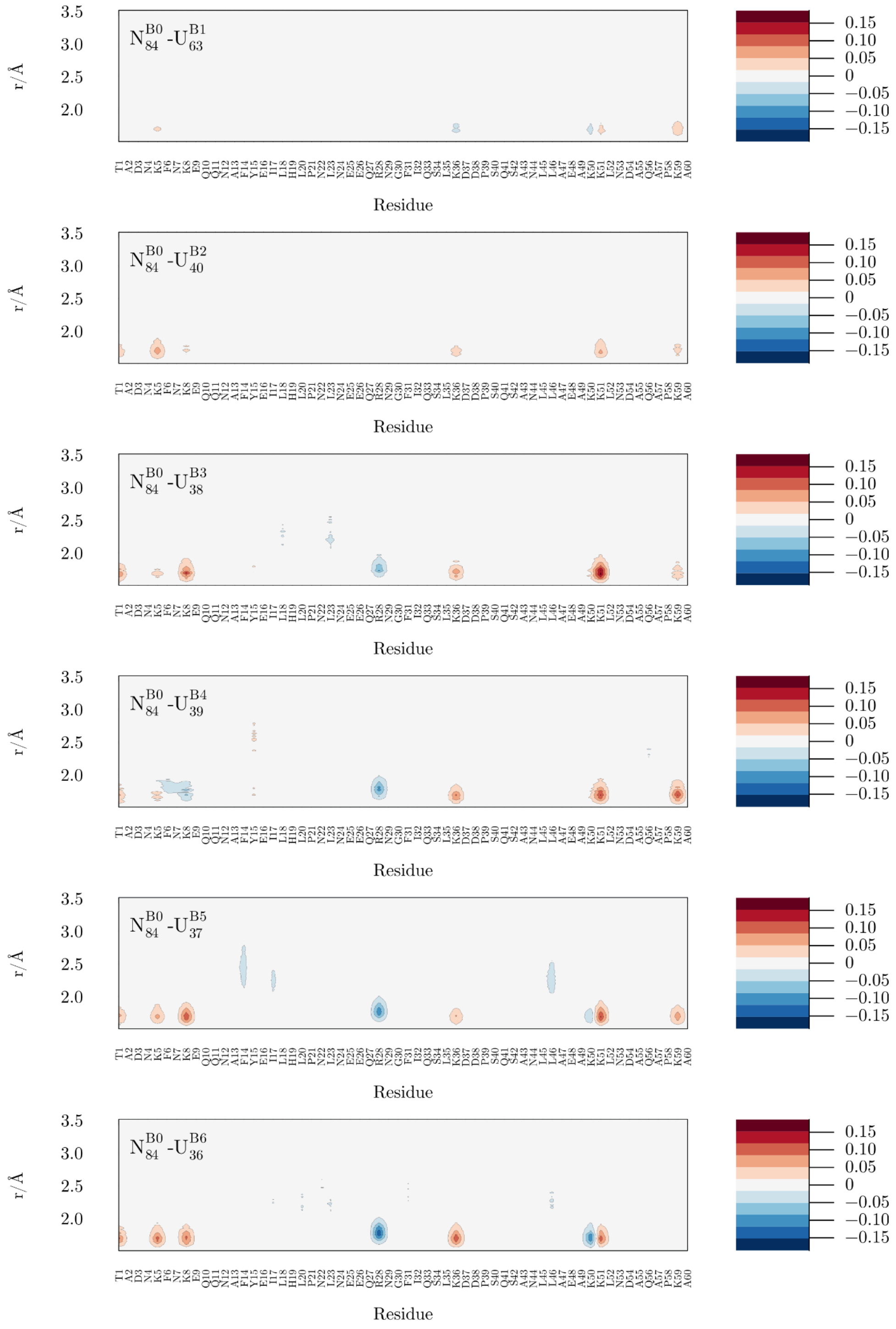


**Figure S13**. Differential density maps per residue for BdpA in TMAO solution 0.5 mol $L^{-1}$.

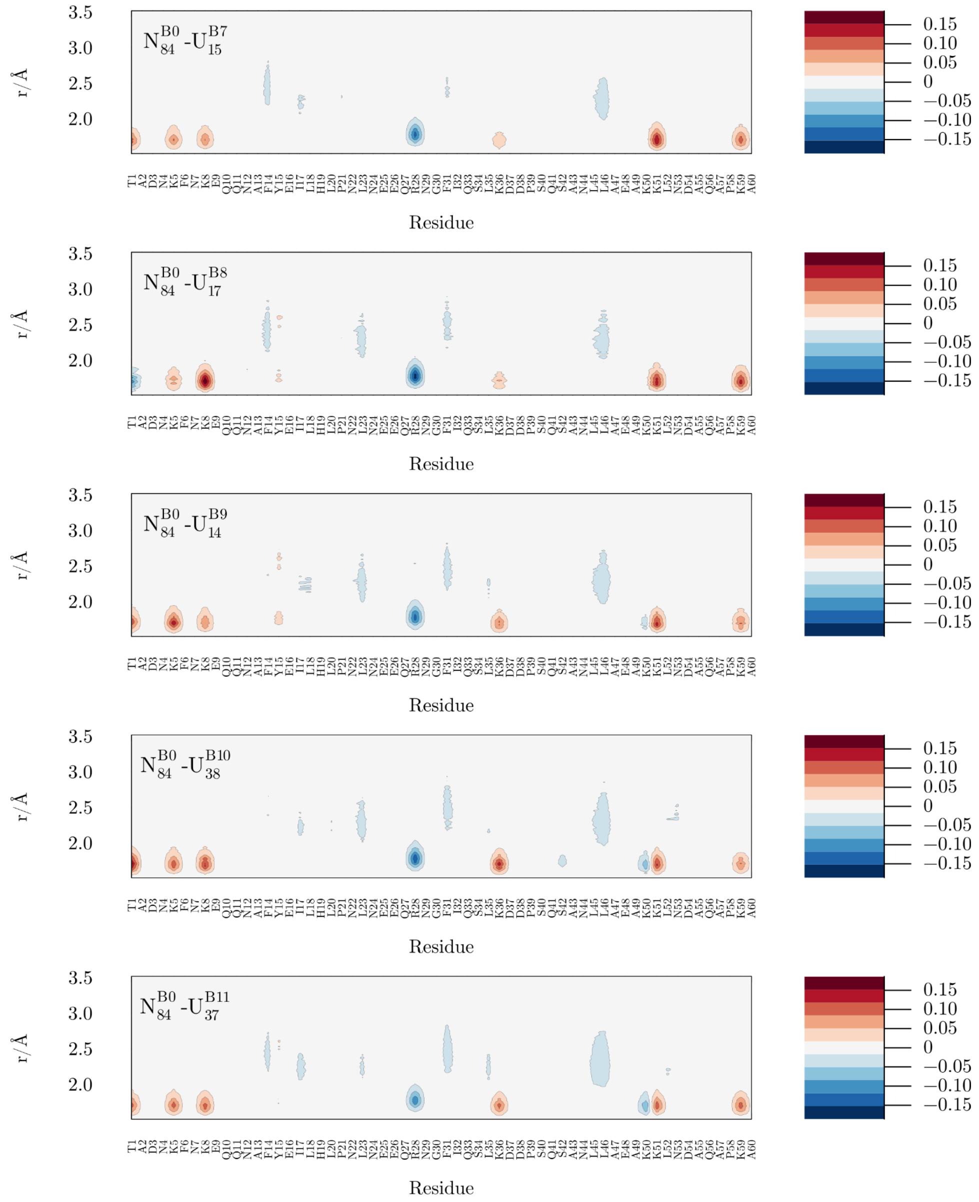


**Figure S13 (continued)**. Differential density maps per residue for BdpA in TMAO solution 0.5 mol $L^{-1}$.

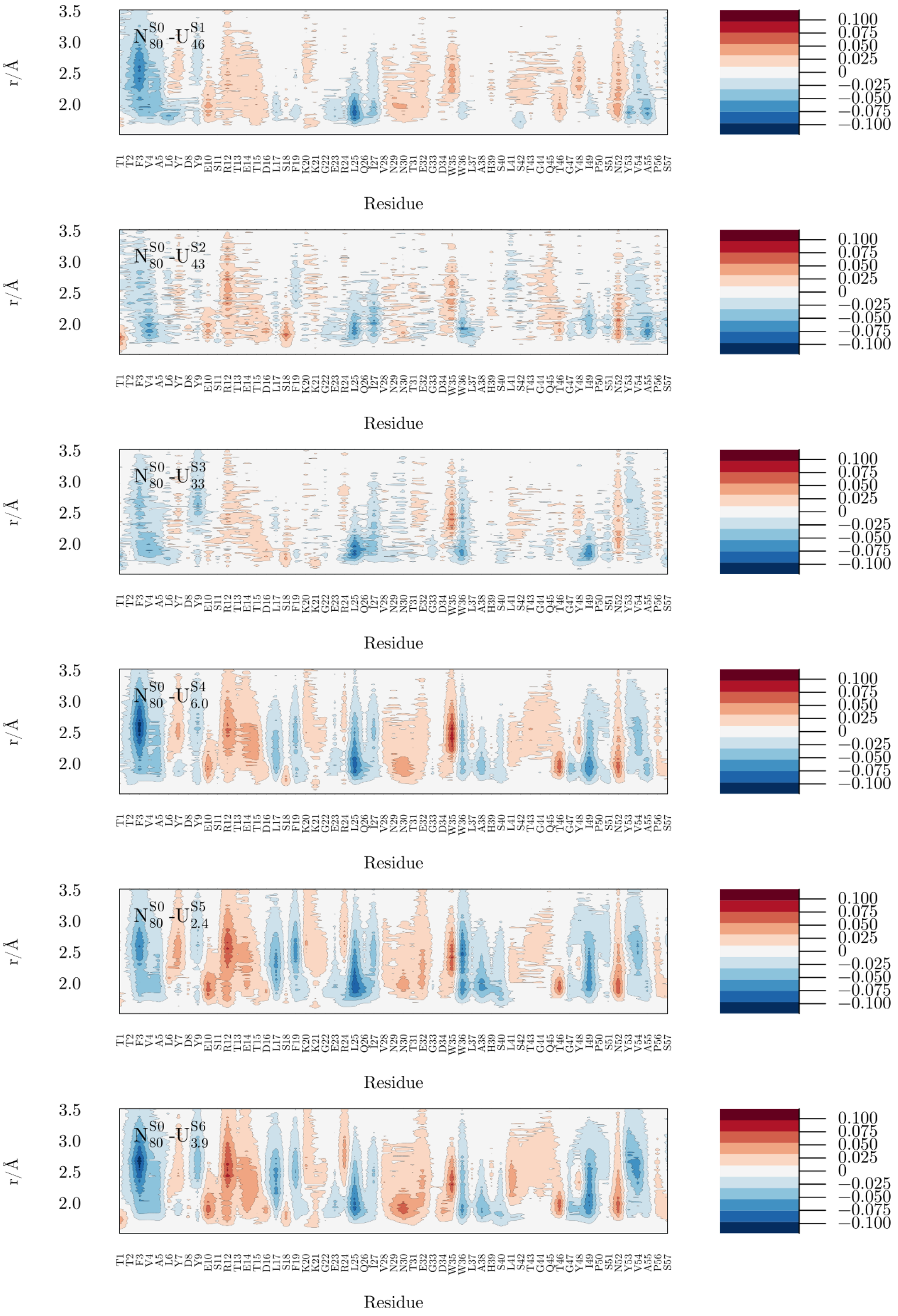


**Figure S14**. Differential density maps per residue for SH3 in urea solution 0.5 mol $L^{-1}$.

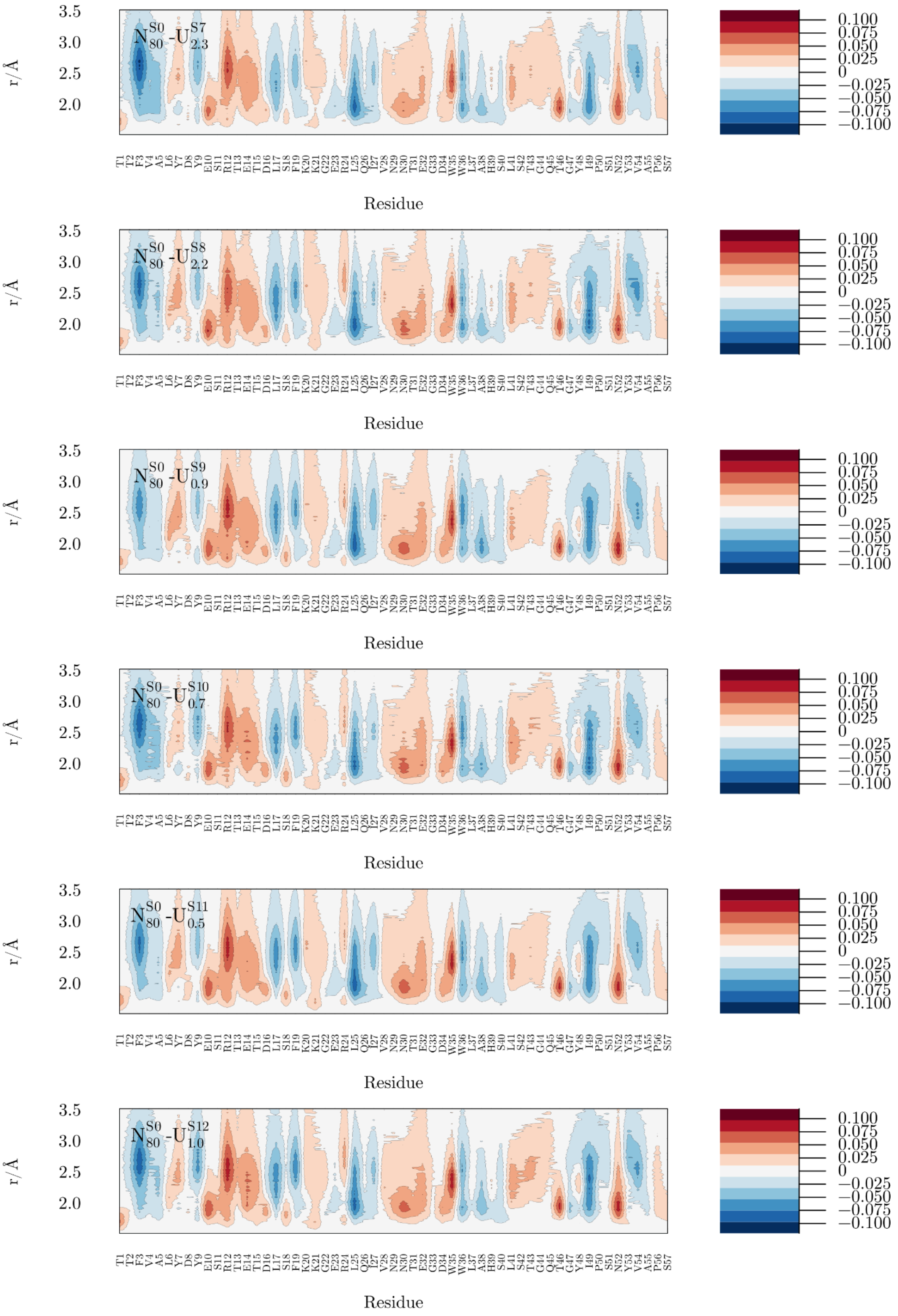


**Figure S14 (continued).** Differential density maps per residue for SH3 in urea solution 0.5 mol $L^{-1}$.

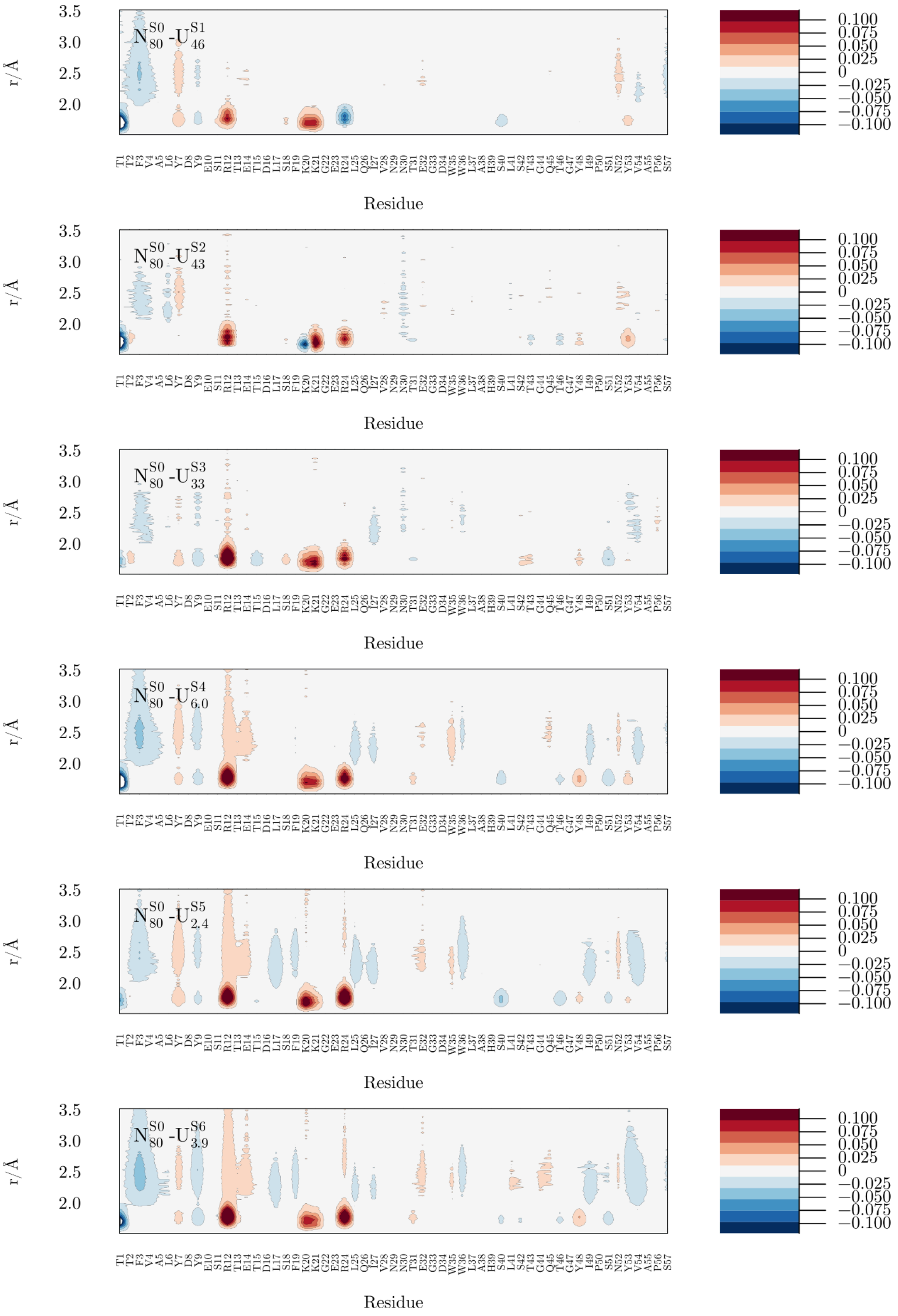


**Figure S15**. Differential density maps per residue for SH3 in TMAO solution 0.5 mol $L^{-1}$.

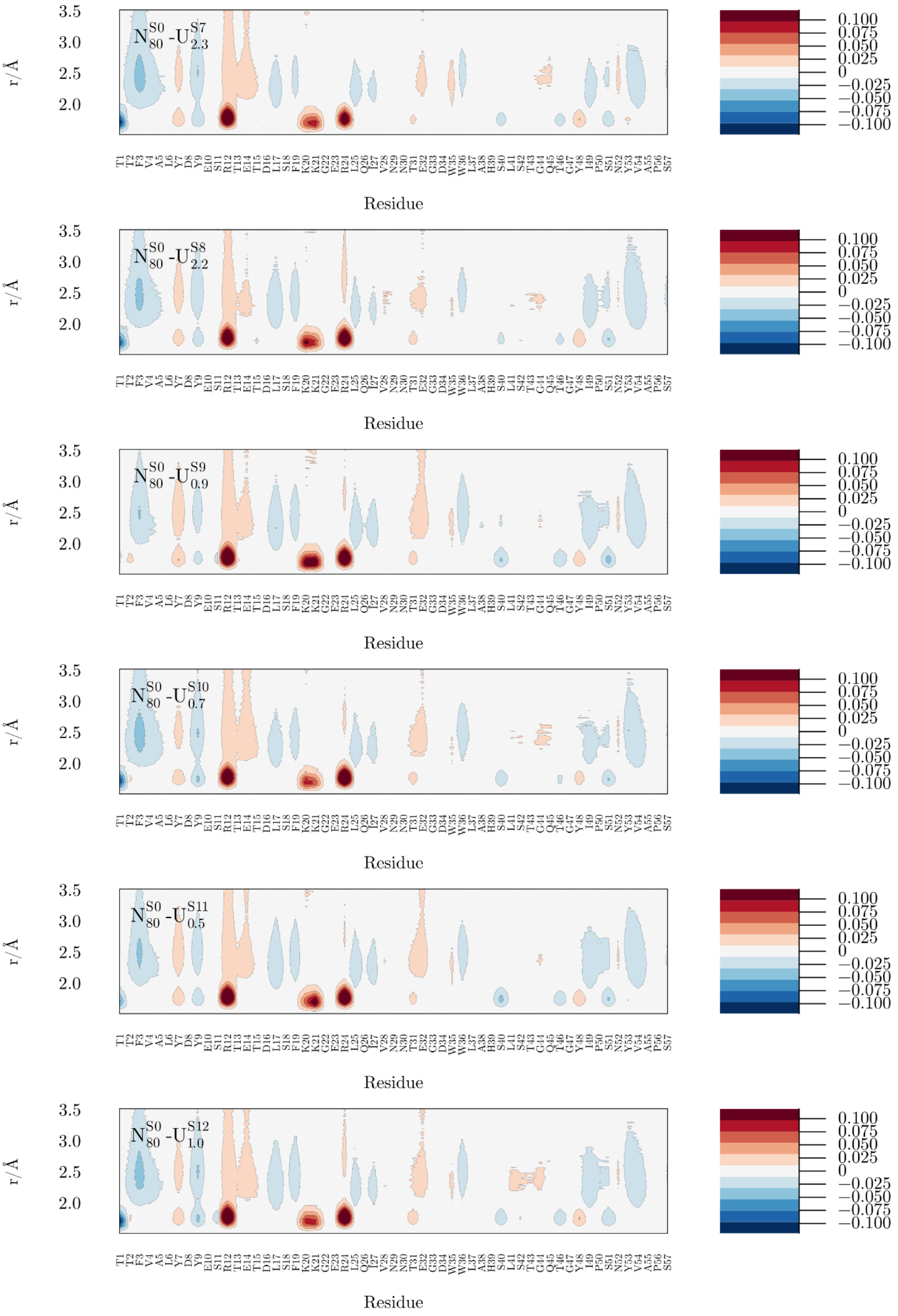


**Figure S15 (continued)**. Differential density maps per residue for SH3 in TMAO solution 0.5 mol L$^{-1}$.

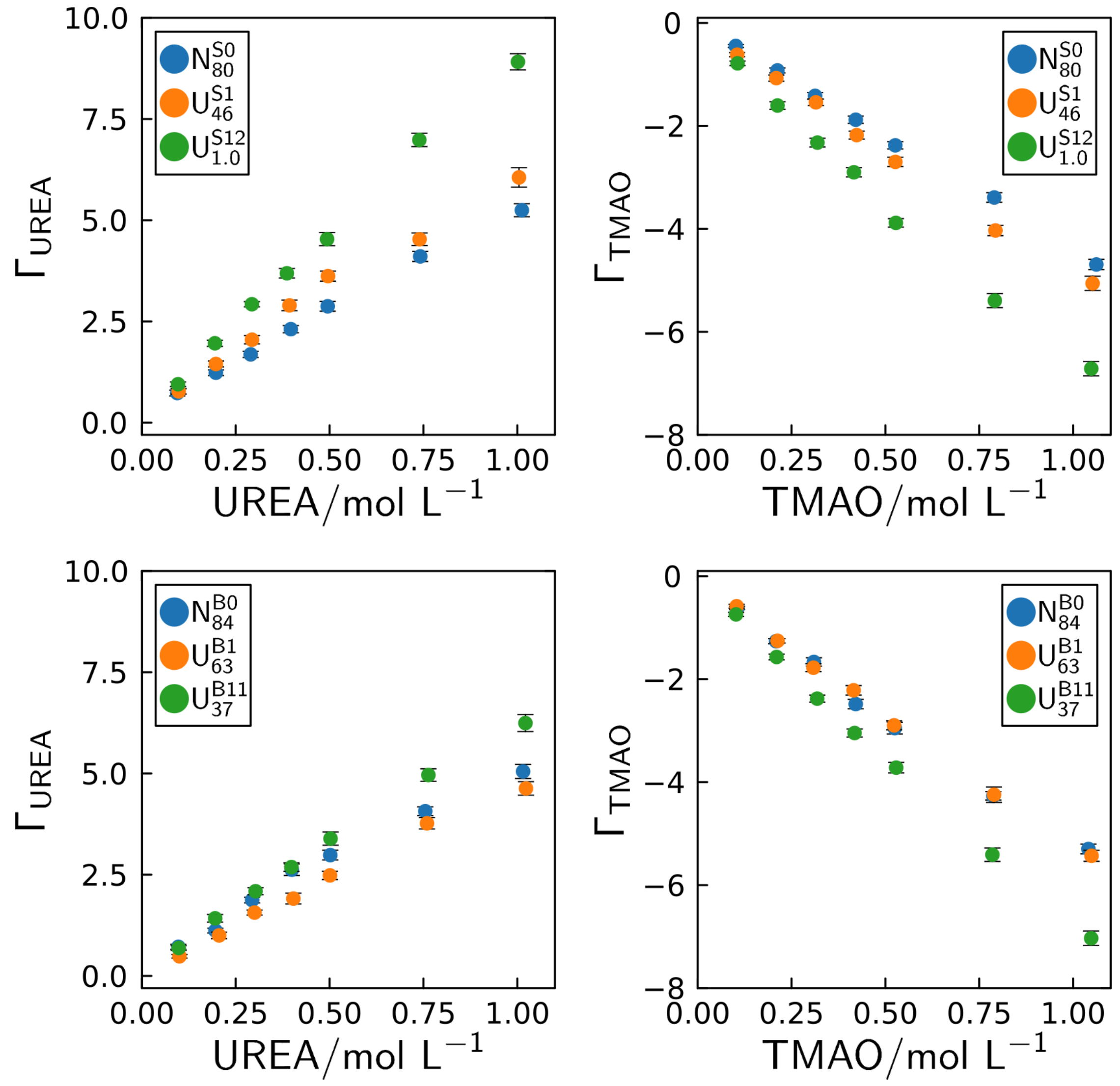


**Figure S16**. Preferential interaction parameters (Γs) as a function of the osmolyte concentrations. A) and B) show the Γ of Urea and TMAO for selected SH3 ensembles, respectively. Similarly, C) and D) show the Γ of Urea and TMAO for selected BdpA ensembles.

**Table S2**. Simulation box details for each SH3 protein ensemble in 0.5 mol L$^{-1}$ urea and TMAO solutions.

| SH3 protein | | | |
|---|---|---|---|
| Ensembles | Average<br>Box Volume (nm3) and SE | Average<br>Water Number and SE | Average<br>Cosolvent Number and SE |
| $N_{80}^{S0}$ | 162.1 ± 0.2 | 4576 ± 6 | 42.78 ± 0.05 |
| $U_{46}^{S1}$ | 182 ± 3 | 5207 ± 86 | 48.7 ± 0.8 |
| $U_{43}^{S2}$ | 180 ± 3 | 5132 ± 82 | 48.0 ± 0.8 |
| $U_{33}^{S3}$ | 177 ± 3 | 5035 ± 86 | 47.1 ± 0.8 |
| $U_{6.0}^{S4}$ | 207 ± 2 | 5957 ± 78 | 55.7 ± 0.7 |
| $U_{2.4}^{S5}$ | 228 ± 3 | 6616 ± 108 | 62 ± 1 |
| $U_{3.9}^{S6}$ | 250 ± 3 | 7296 ± 101 | 68.2 ± 0.9 |
| $U_{2.3}^{S7}$ | 249 ± 3 | 7268 ± 82 | 67.9 ± 0.8 |
| $U_{2.2}^{S8}$ | 284 ± 4 | 8353 ± 129 | 78 ± 1 |
| $U_{0.9}^{S9}$ | 330 ± 7 | 9806 ± 234 | 92 ± 2 |
| $U_{0.7}^{S10}$ | 282 ± 4 | 8286 ± 113 | 78 ± 1 |
| $U_{0.5}^{S11}$ | 322 ± 4 | 9549 ± 112 | 89 ± 1 |
| $U_{1.0}^{S12}$ | 313 ± 4 | 9268 ± 124 | 87 ± 1 |

**Table S3**. Simulation box details for each BdpA protein ensemble in 0.5 mol $L^{-1}$ urea and TMAO solutions.

| BdpA protein | | | |
|---|---|---|---|
| Ensembles | Average<br>Box Volume (nm3) and SE | Average<br>Water Number and SE | Average<br>Cosolvent Number and SE |
| $N_{84}^{B0}$ | 208.1 ± 0.9 | 5985 ± 28 | 55.9 ± 0.3 |
| $U_{63}^{B1}$ | 207.1 ± 0.8 | 5953 ± 23 | 55.7 ± 0.2 |
| $U_{40}^{B2}$ | 203 ± 1 | 5822 ± 37 | 54.4 ± 0.3 |
| $U_{38}^{B3}$ | 191 ± 2 | 5467 ± 68 | 51.1 ± 0.6 |
| $U_{39}^{B4}$ | 192 ± 3 | 5496 ± 94 | 51.2 ± 0.9 |
| $U_{37}^{B5}$ | 242 ± 2 | 7023 ± 55 | 65.7 ± 0.5 |
| $U_{36}^{B6}$ | 221 ± 2 | 6392 ± 72 | 59.8 ± 0.7 |
| $U_{15}^{B7}$ | 235 ± 2 | 6818 ± 74 | 63.7 ± 0.7 |
| $U_{17}^{B8}$ | 251 ± 4 | 7324 ± 120 | 68 ± 1 |
| $U_{14}^{B9}$ | 223 ± 3 | 6446 ± 104 | 60 ± 1 |
| $U_{38}^{B10}$ | 240 ± 4 | 6973 ± 110 | 65 ± 1 |
| $U_{37}^{B11}$ | 302 ± 4 | 8909 ± 105 | 83 ± 1 |

**Table S4**. Simulation box details for each representative SH3 protein structure in 0.1, 0.2, 0.3, 0.4, 0.5, 0.75, 1.0 mol L$^{-1}$ urea and TMAO solutions.

| SH3 protein | | | | |
|---|---|---|---|---|
| Representative structure | Concentration (mol L$^{-1}$) | Box Volume (nm$^3$) | Water Number | Cosolvent Number |
| $N_{80}^{S0}$ | 0.1 | 228.54 | 6826 | 12 |
| $N_{80}^{S0}$ | 0.2 | 227.73 | 6772 | 25 |
| $N_{80}^{S0}$ | 0.3 | 227.79 | 6722 | 37 |
| $N_{80}^{S0}$ | 0.4 | 227.60 | 6667 | 50 |
| $N_{80}^{S0}$ | 0.5 | 227.71 | 6617 | 62 |
| $N_{80}^{S0}$ | 0.75 | 228.00 | 6488 | 93 |
| $N_{80}^{S0}$ | 1 | 228.11 | 6359 | 124 |
| $U_{46}^{S1}$ | 0.1 | 254.66 | 7674 | 14 |
| $U_{46}^{S1}$ | 0.2 | 255.27 | 7615 | 28 |
| $U_{46}^{S1}$ | 0.3 | 254.78 | 7557 | 42 |
| $U_{46}^{S1}$ | 0.4 | 255.14 | 7499 | 56 |
| $U_{46}^{S1}$ | 0.5 | 255.04 | 7440 | 70 |
| $U_{46}^{S1}$ | 0.75 | 255.39 | 7298 | 104 |
| $U_{46}^{S1}$ | 1 | 254.65 | 7152 | 139 |
| $U_{1.0}^{S12}$ | 0.1 | 322.87 | 9851 | 18 |
| $U_{1.0}^{S12}$ | 0.2 | 322.07 | 9776 | 36 |
| $U_{1.0}^{S12}$ | 0.3 | 323.07 | 9701 | 54 |
| $U_{1.0}^{S12}$ | 0.4 | 322.02 | 9630 | 71 |
| $U_{1.0}^{S12}$ | 0.5 | 322.17 | 9555 | 89 |
| $U_{1.0}^{S12}$ | 0.75 | 322.43 | 9367 | 134 |
| $U_{1.0}^{S12}$ | 1 | 322.49 | 9179 | 179 |

**Table S5**. Simulation box details for each representative BdpA protein structure in 0.1, 0.2, 0.3, 0.4, 0.5, 0.75, 1.0 mol L$^{-1}$ urea and TMAO solutions.

| BdpA protein | | | | |
|---|---|---|---|---|
| Representative structure | Concentration (mol L$^{-1}$) | Box Volume (nm$^3$) | Water Number | Cosolvent Number |
| $N_{84}^{B0}$ | 0.1 | 327.229 | 10000 | 18 |
| $N_{84}^{B0}$ | 0.2 | 327.344 | 9924 | 36 |
| $N_{84}^{B0}$ | 0.3 | 327.15 | 9849 | 54 |
| $N_{84}^{B0}$ | 0.4 | 327.72 | 9770 | 73 |
| $N_{84}^{B0}$ | 0.5 | 327.294 | 9695 | 91 |
| $N_{84}^{B0}$ | 0.75 | 327.065 | 9507 | 136 |
| $N_{84}^{B0}$ | 1 | 327.14 | 9319 | 181 |
| $U_{63}^{B1}$ | 0.1 | 305.938 | 9262 | 17 |
| $U_{63}^{B1}$ | 0.2 | 306.058 | 9191 | 34 |
| $U_{63}^{B1}$ | 0.3 | 306.046 | 9124 | 50 |
| $U_{63}^{B1}$ | 0.4 | 305.645 | 9053 | 67 |
| $U_{63}^{B1}$ | 0.5 | 306.011 | 8983 | 84 |
| $U_{63}^{B1}$ | 0.75 | 306.166 | 8807 | 126 |
| $U_{63}^{B1}$ | 1 | 305.406 | 8632 | 168 |
| $U_{37}^{B11}$ | 0.1 | 396.974 | 11673 | 21 |
| $U_{37}^{B11}$ | 0.2 | 397.009 | 11585 | 42 |
| $U_{37}^{B11}$ | 0.3 | 396.789 | 11493 | 64 |
| $U_{37}^{B11}$ | 0.4 | 398.107 | 11406 | 85 |
| $U_{37}^{B11}$ | 0.5 | 396.861 | 11318 | 106 |
| $U_{37}^{B11}$ | 0.75 | 397.379 | 11097 | 159 |
| $U_{37}^{B11}$ | 1 | 397.272 | 10876 | 212 |
